# The Role of ATP Synthase Subunit e (ATP5I) in Mediating the Metabolic and Antiproliferative Effects of Metformin

**DOI:** 10.1101/2024.09.20.614047

**Authors:** Guillaume Lefrançois, Emilie Lavallée, Marie-Camille Rowell, Véronique Bourdeau, Farzaneh Mohebali, Thierry Bertomeu, Ana Maria Duman, Maya Nikolova, Mike Tyers, Simon-Pierre Gravel, Andréea R. Schmitzer, Gerardo Ferbeyre

## Abstract

Here we identify the subunit e of F F -ATP synthase (ATP5I) as a target of metformin, a first-in-class antidiabetic biguanide. ATP5I maintains the stability of F F -ATP synthase dimers which is crucial for shaping cristae morphology. We demonstrate that ATP5I interacts with a biguanide analogue *in vitro* and disabling its expression by CRISPR-Cas9 in pancreatic cancer cells leads to the same phenotype as biguanide treated cells including mitochondrial morphology alterations, reduction of the NAD^+^/NADH ratio, inhibition of oxidative phosphorylation (OXPHOS), rescue of respiration by uncouplers and a compensatory increase in glycolysis. Notably, metformin affects the oligomerization of the F F -ATP synthase leading to accumulation of vestigial assembly intermediates also observed upon ATP5I inactivation. Moreover, ATP5I knockout (KO) cells exhibit resistance to the antiproliferative effects of biguanides, but reintroduction of ATP5I rescues the metabolic and anti-proliferative effects of metformin and phenformin. Finally, a genome-wide CRISPR screening in NALM-6 lymphoma cells revealed that metformin-treated cells exhibit genetic interaction profiles similar to those observed with the F F -ATP synthase inhibitor oligomycin, but not with the complex I inhibitor rotenone. This provides unbiased support for the relevance of the newly proposed target.

## Introduction

Discovering safe and effective therapeutic targets for cancer treatment remains a major challenge in biomedical research^1^. Recently, targeted therapies against complex I of the respiratory chain have garnered considerable attention, offering a promising strategy for cancer treatment due to their potential to disrupt the energy metabolism of tumor cells^2–5^. However, despite initial promises, clinical trials with potent complex I inhibitors have been compromised by severe toxicities, raising concerns about their clinical viability. Given the critical role of mitochondria in cancer cells, there is an urgent need for more effective and safer alternatives to target them^6,7^. In this context, medicinal biguanides such as metformin and the more lipophilic phenformin are emerging as attractive alternatives. These biguanides are considered moderate respiratory chain inhibitors and offer a potential avenue for targeting mitochondrial metabolism in cancer^8^.

Metformin has been used since the 1950s for its anti-hyperglycemic properties^9^. As such, it has a well-established safety profile and is commonly used for the treatment of type II diabetes^10^. Several epidemiological studies showed that prolonged use of metformin in diabetic patients reduced the incidence of several cancers^11,12^, particularly pancreatic cancer^13,14^. However, its clinical use has proven ineffective in patients with advanced pancreatic cancer^15^, suggesting that its antitumoral mechanism is more preventive than therapeutic^16,17^. The more potent phenformin^9,^ which was withdrawn from the market in the 1970s for its anti-hyperglycemic applications due to deaths attributed partly to lactic acidosis^18^, is currently undergoing clinical trials in cancer patients ^19^.

Unlike other inhibitors of the respiratory chain, the mechanism of action of medicinal biguanides remains elusive^20,21^. However, compelling evidence indicates that energy metabolism plays a pivotal role in their effectiveness^22,23^. At supra-pharmacological doses (1-10 mM), typically 100-1000 times higher doses than those used for anti-hyperglycaemic effects, metformin moderately inhibits the activity of complex I of the respiratory chain in several cellular models^24–26^. This inhibition subsequently decreases OXPHOS activity, leading to reduced cellular respiration and energy production. In response, cells adapt by shifting metabolism to glycolysis and decreasing their anabolic activity through the activation of the energy sensor AMPK^20,22,24^. Nonetheless, to observe a decrease in isolated complex I activity in *in vitro* models, metformin doses of up to 50 mM are required^25–27^. A recent cryo-electron microscopy study revealed that a metformin analogue binds to complex I in a specific conformation with a binding site unique to this analogue^27^. However, further work using *in vivo* models is necessary to validate the biological meaning of this interaction as a critical site of action of metformin^27,28^.

Additional targets such as glycerol phosphate dehydrogenase^29^, complex IV^30^, PEN2^31^ and F F -ATP synthase^25^ have also been suggested for metformin. Metformin may interact with various targets in a nonspecific manner^32–35^ because of its structural similarity with guanidine, a potent chaotropic agent. These various targets potentially exert pleiotropic effects^20,21^ that could act synergistically.

It has been shown that atrazine, a molecule derived from biguanides, binds and inhibits F F - ATP synthase^36^. Additionally, a study from our group revealed that medicinal biguanides likely disrupt cristae organization^37^, a crucial process mediated by the oligomerization of F F -ATP synthase and resulting in the formation of ⍰⍰onion-liked structures⍰⍰^38–41^. It is proposed that complex I and F F -ATP synthase of the respiratory chain may be particularly sensitive to biguanide inhibition due to the conformational mobility of their catalytic interfaces^25^. Disrupting the folding of the mitochondrial inner membrane could alter the organization and functions of other respiratory chain complexes^42^. It remains an open question whether biguanides act through multiple targets or a single primary target.

Here, we identify another potential target of biguanides: the e subunit of F F -ATP synthase (ATP5I), a transmembrane protein involved in the dimerization and assembly of this supramolecular complex^38–40^. Although medicinal biguanides are known to accumulate in mitochondria and disrupt the respiratory chain by inhibiting complex I, no direct evidence of their interaction with a subunit of F F -ATP synthase has been reported until now. CRISPR-Cas9 mediated KO of ATP5I recapitulates many of the effects of metformin in pancreatic cancer cells and reduces metformin sensitivity. Conversely, treatment with metformin interferes with the oligomerization and assembly of the F F -ATP synthase complex as happens in cells without ATP5I. Interestingly, the genetic signature of cells treated with metformin is more similar to that of cells treated with an F F -ATP synthase inhibitor than to a complex I inhibitor. Taken together this work adds the F F -ATP synthase and its subunit ATP5I as a bona fide target of biguanides.

## Results

### Identification of ATP synthase subunit e (ATP5I) as a mitochondrial biguanide binding protein

To identify proteins that bind biguanides, we synthetized a Biotin Functionalized Biguanide or BFB (Figure 1A, Figure 1 – figure supplement 1A-B) to perform an affinity-based pull-down assay with streptavidin beads^43^. We first assessed the biological activity of BFB comparing it to metformin in the ability to activate AMPK and inhibit cell proliferation in KP-4 pancreatic cancer cells. BFB activated AMPK and its downstream target ACC as indicated by their increased phosphorylation states (Figure 1B) and inhibited KP-4 cell growth with an EC_50_ of 1.0 ± 0.2 mM similar to metformin (Figure 1C). Additionally, immunofluorescence experiments with fluorophore-conjugated Streptavidin confirmed that BFB accumulates in mitochondria, as shown by colocalization with the mitochondrial protein TOMM20 (Figures 1D, 1E).

**Figure 1.**
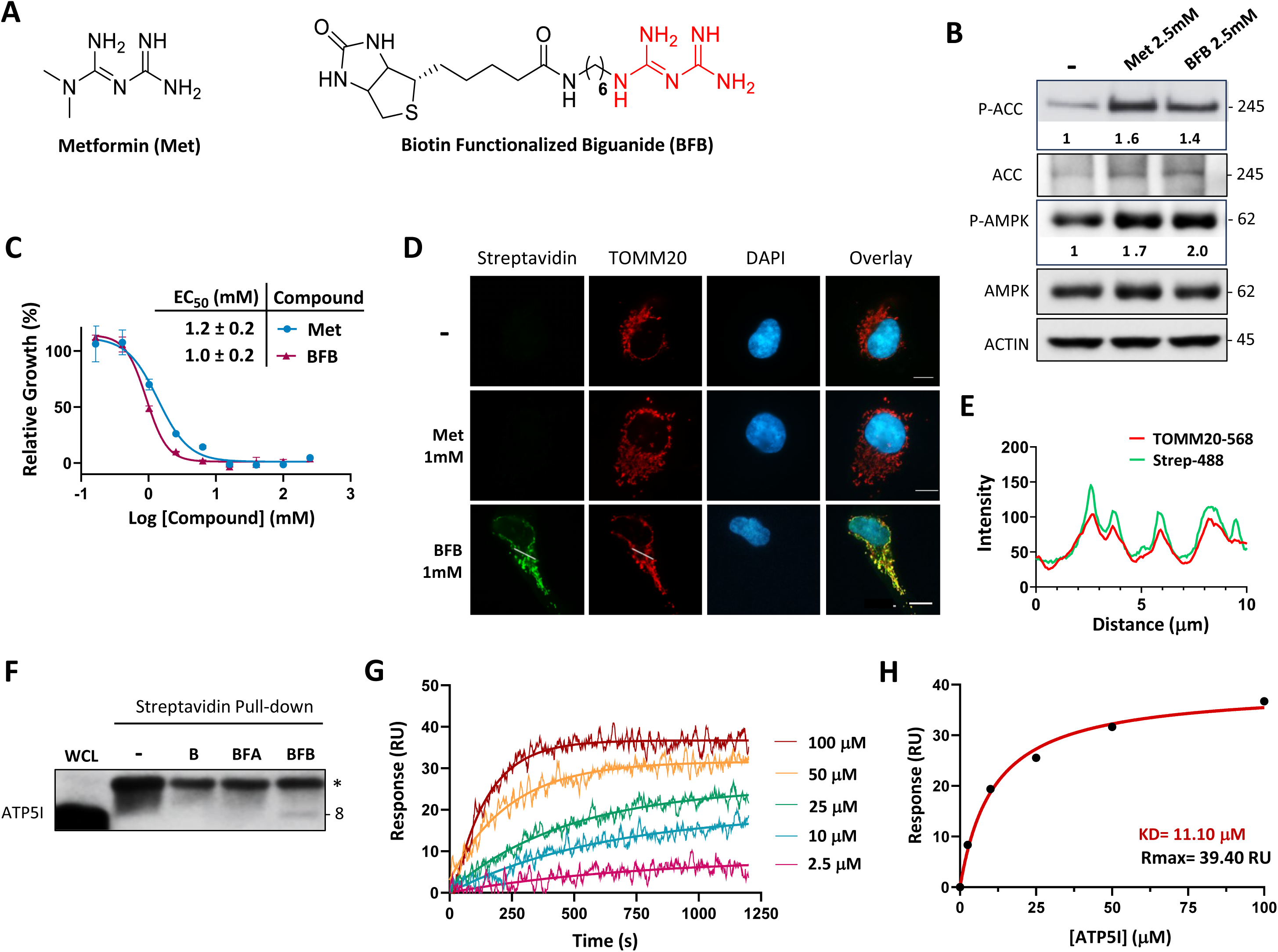
Biguanide pharmacophore interacts with ATP synthase subunit e (ATP5I). **(A)** Design of bio-inspired probe biotin functionalized biguanide (BFB) based on the structure of metformin (Met). **(B)** Immunoblots for the phosphorylation of AMPK (Thr172) and ACC (Ser79) in extracts from KP-4 pancreatic cancer cells treated with 2.5 mM Met or BFB for 16 hours. β-ACTIN was used as loading control. **(C)** Representative quantification of cell viability and growth with corresponding EC_50_ values of 3-day treatments with metformin (Met) or biotin functionalized biguanide (BFB) in KP-4 cells. Values represent the mean ± standard deviation of N=3. **(D)** Representative images of mitochondria and BFB localization in cells as in (B). Cells were treated with 1 mM of metformin (Met) or BFB for 16 hours and mitochondrial signal and BFB localization were analyzed by co-immunofluorescence using Streptavidin fluorophore conjugate and anti-TOMM20 antibody, scale bar= 10 μm. Cells untreated (-) and treated with 1 mM Met were used as negative controls. **(E)** Colocalization between TOMM20 (TOMM20-568) and Streptavidin (Strep-488) fluorophores was analyzed for the BFB condition from (D) through job plot intensity profile. **(F)** Pull-down validation experiments with streptavidin beads alone (-), D-biotin (B), BFA and BFB using antibody followed by immunoblot against ATP5I in cells as in in extracts from HEK-293T embryonic kidney cells. The whole cell lysate (WCL) was added as control. **(G)** Binding interactions studies of BFB with recombinant purified ATP5I (rATP5I) using Surface Plasmons of Resonance (SPR). Representative sensorgrams show affinity kinetics of BFB and rATP5I. BFB was exposed onto streptavidin immobilized sensor chip and several concentrations of rATP5I were added until saturation of the signal. RU: Resonance Units. **(H)** Binding affinity curve obtained from each steady state from (F). K_D_ refers to the dissociation equilibrium constant and Rmax represent the theoretical maximum response.

**Figure 1 – figure supplement 1.**
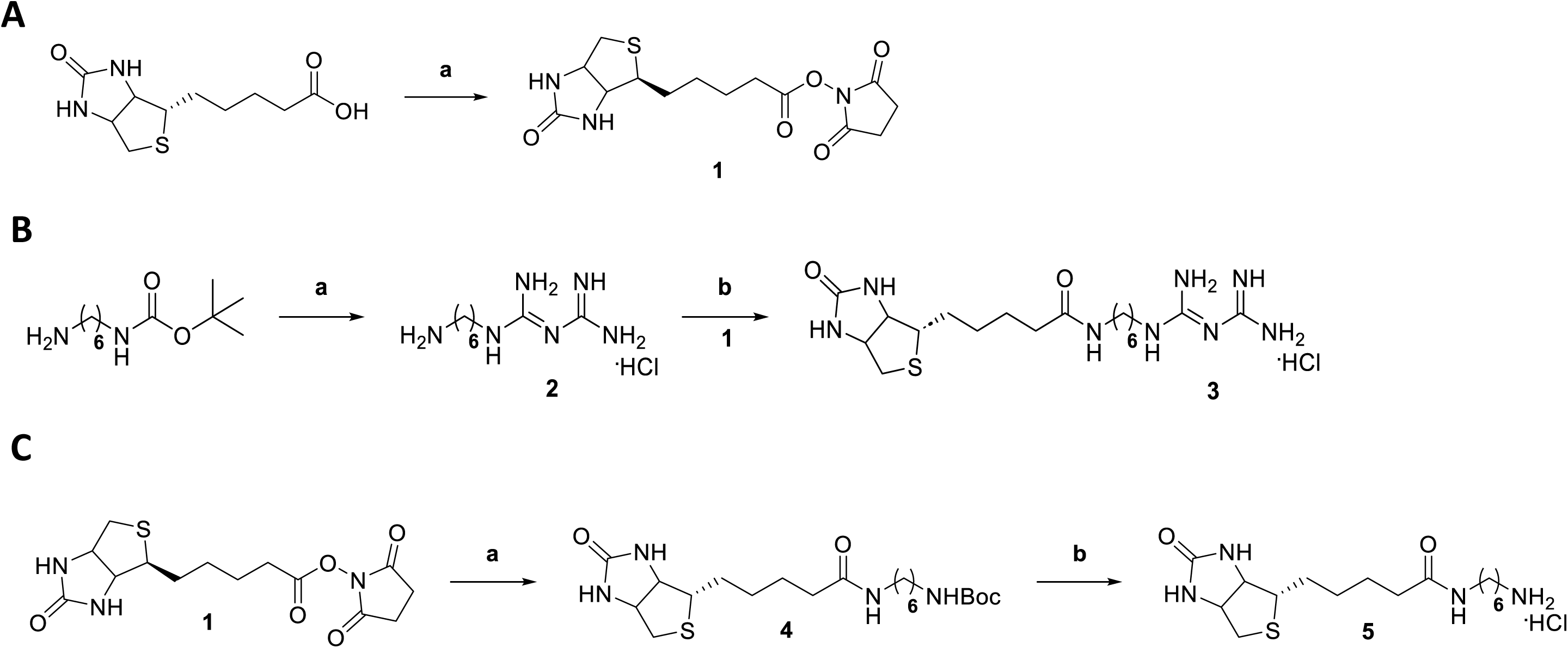
**(A)** Synthesis of biotin-NHS (1); (a) NHS, EDC, DMF, room temperature (r.t) overnight (o/n). **(B)** Synthesis of biotin functionalized biguanide (BFB) chloride salt (3); (a) Dicyandiamide, TMSCl, MeCN, 160°C, 3h, 2: 6-aminohexylbiguanide hydrochloride salt; (b) Biotin-NHS, DIPEA, DMF, r.t, o/n. **(C)** Synthesis of biotin functionalized amine (BFA) chloride salt (5); (a) N-Boc-1,6-hexanediamine, DMF, r.t, o/n, 4: biotin functionalized N-Boc-amine; (b) HCl/MeOH, MeOH, r.t, o/n.

**Figure 1 – figure supplement 2.**
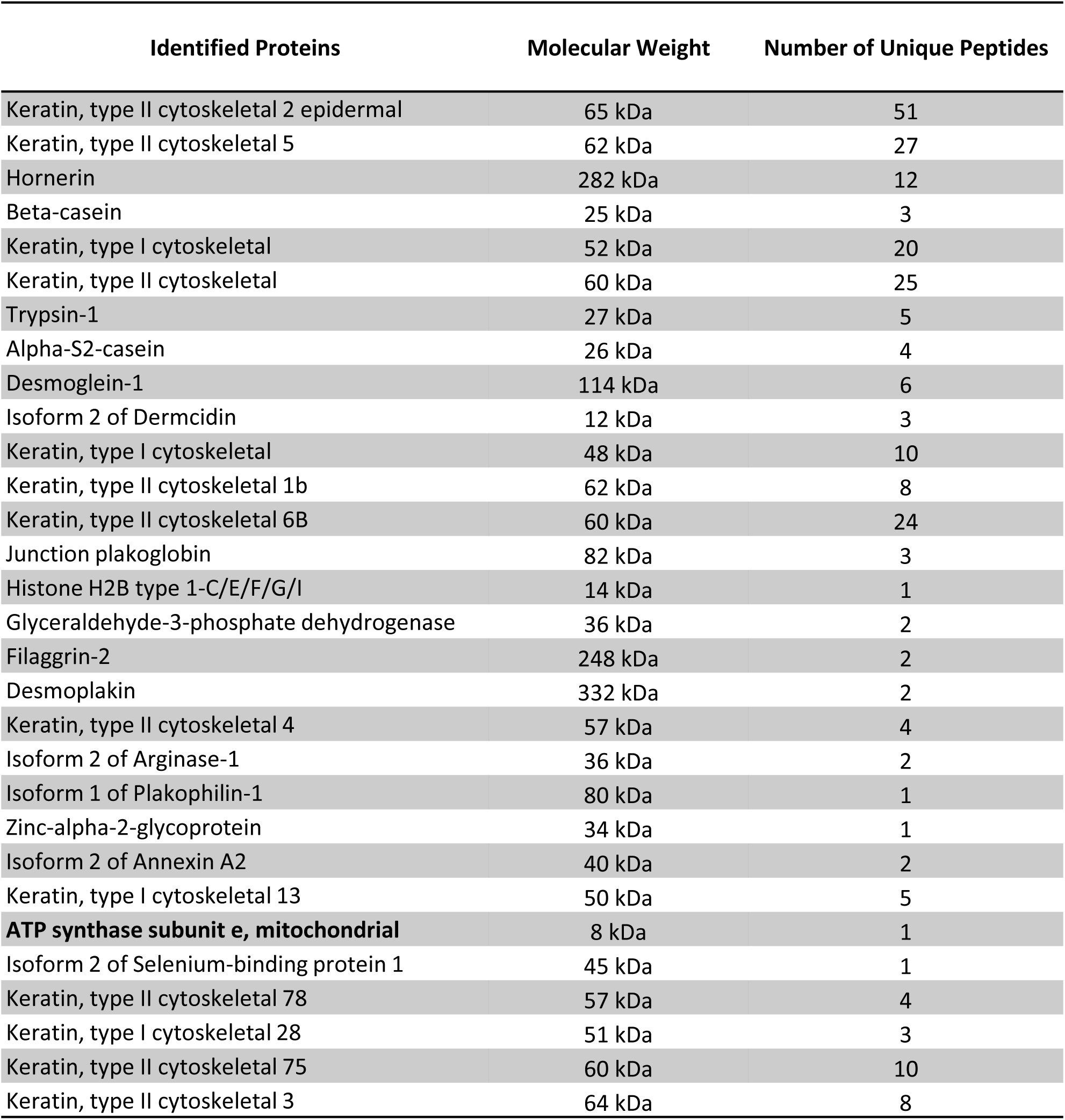
Proteins were isolated from streptavidin-coated beads after affinity purification using the biotinylated biguanide probe (BFB), followed by competitive elution with metformin (50 mM). Mass spectrometry (MS/MS) analysis was performed to identify proteins potentially involved in BFB-specific interactions. The table reports the approximate molecular weight and total number of peptides identified for each protein.

**Figure 1 – figure supplement 3.**
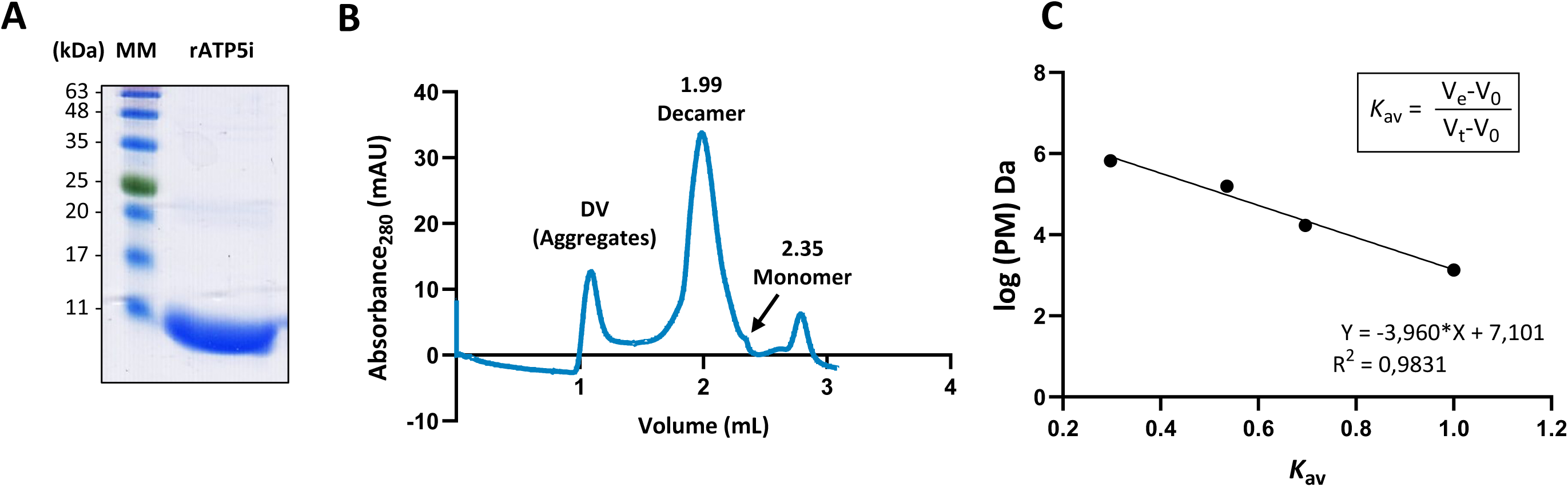
**(A)** SDS PAGE analysis of recombinant ATP5I (rATP5I) after the purification process. Purity is estimated at ≥ 90 %. MM: molecular weight marker. **(B)** Gel filtration profile analysis of rATP5I using Superose 12 10/300 GL. DV represents column dead volume. mAU: milli-Absorbance Unit. **(C)** Calibration of the Superose 6 10/300 GL size-exclusion column for conformational analysis of the Nter-6×His-ATP5I construct. The calibration curve was generated by plotting the logarithm of the molecular weight (log(MW)) of standard proteins (thyroglobulin, γ-globulin, myoglobin, vitamin B12) as a function of their partition coefficient (Kav). Ve: elution volume; V : void volume (1.12 mL); Vt: total column volume (2.8 mL).

After confirming that BFB exhibited activity comparable to that of metformin, we performed pull-down assays using streptavidin immobilized on sepharose beads on mitochondrial-enriched cell extracts to analyse interacting proteins by mass-spectrometry. Given the significant issue of nonspecific binding associated with this technique, we performed multiple parallel pull-down experiments. These included a control biotin-conjugated amine derivative (BFA) possessing the same linker as BFB (Figure 1 – figure supplement 1C). Mass spectrometry analysis results identified a total of 69 proteins. Among these, 30 proteins showed specific interaction with BFB under the elution condition with metformin (Figure 1 – figure supplement 2). Keratins identified in this way were considered contaminants. However, two mitochondrial proteins stood out, arginase isoform 2 and ATP synthase subunit e (ATP5I). Arginase binds arginine which has a guanidinium group structurally related to metformin^44^ while ATP5I is part of the peripheral stalk of the F F -ATP synthase ^45^.

### Biguanide pharmacophore interacts specifically with ATP5I

As ATP5I was not previously shown to bind biguanides, we first confirmed this interaction through immunoblot analysis in independent pull-downs experiments. The results indicate that BFB allows specific interaction of ATP5I in streptavidin pull-down while BFA failed to do so. Of note, we observed a nonspecific band (≃11 kDA) in all pull-down conditions with streptavidin beads, possibly associated with the denatured streptavidin monomer in heated SDS buffer (Figure 1F). We then used surface plasmon resonance (SPR) to characterize the binding of BFB and purified recombinant ATP5I. Our purification steps yielded a highly pure recombinant protein, mainly organized as oligomers (mostly decamers) in solution (Figure 1– figure supplement 3A-C). Although SPR is sensitive^46,47^, the size difference between BFB and recombinant ATP5I prompted us to immobilize the small molecule BFB on the gold surface. Adding increasing concentrations of recombinant ATP5I allowed us to estimate an affinity constant (KD ≃ 11.10 μM; Rmax ≃ 39.40 RU) significantly lower than typical metformin concentrations that affect complex I in mM range (Figures 1G, 1H).

### ATP5I knockout in pancreatic cancer cells alters the organization of mitochondrial networks

In yeast, inhibition of the subunit e’s equivalent impaired mitochondrial inner membrane folding^48^. This structural deficiency in yeast correlates with decreased dimerization of F F - ATP synthase, highlighting the essential role of ATP5I in dimer stability. However, yeast’s subunit e is not essential for the catalytic activity of F F -ATP synthase^48–52^. In human cells, ATP5I is also crucial for dimer stability^40^ but lacks thorough characterization regarding its roles in cellular energy metabolism. To elucidate its function in cancer and its relevance to cells treated with medicinal biguanides, we developed CRISPR-based reagents to inhibit ATP5I expression in KP-4 pancreatic cancer cells.

Initially, two clones each of the two guide (sgATP5I #1 and sgATP5I #2) were isolated, and their mitochondrial phenotype characterized. First, we measured levels of several proteins from the F F -ATP synthase and respiratory complexes. The results show that all ATP5I knockout clones (ATP5I KO) did not express ATP5I or its partner ATP5L (subunit g in yeast) and had lower levels of ATP5O (oligomycin sensitivity-conferring protein, OSCP in yeast) (Figure 2A). This result is consistent with previous data in yeast where knockout of subunit e also led to a decrease in subunit g ^52^. ATP5I KO cells also exhibited decreased expression of complex I NDUFB8 protein and complex IV COX II protein compared to control clones expressing a guide against GFP (sgGFP) (Figure 2A and Figure 2– figure supplement 1A). A moderate to no reduction in mRNA levels encoding these proteins were observed (Figure 2 – figure supplement 1B) cannot account for the downregulations at the protein level. Additionally, the mitochondrial/nuclear DNA ratio (Figure 2 – figure supplement 1C) indicates no reduction in mitochondrial number, suggesting ATP5I may play a role in maintaining the stability of certain subunits of complexes I and IV. Consistent with this finding, the absence of subunit e in yeast also controls the stability of F F -ATPase proteins subunit g and subunit k^52^.

**Figure 2.**
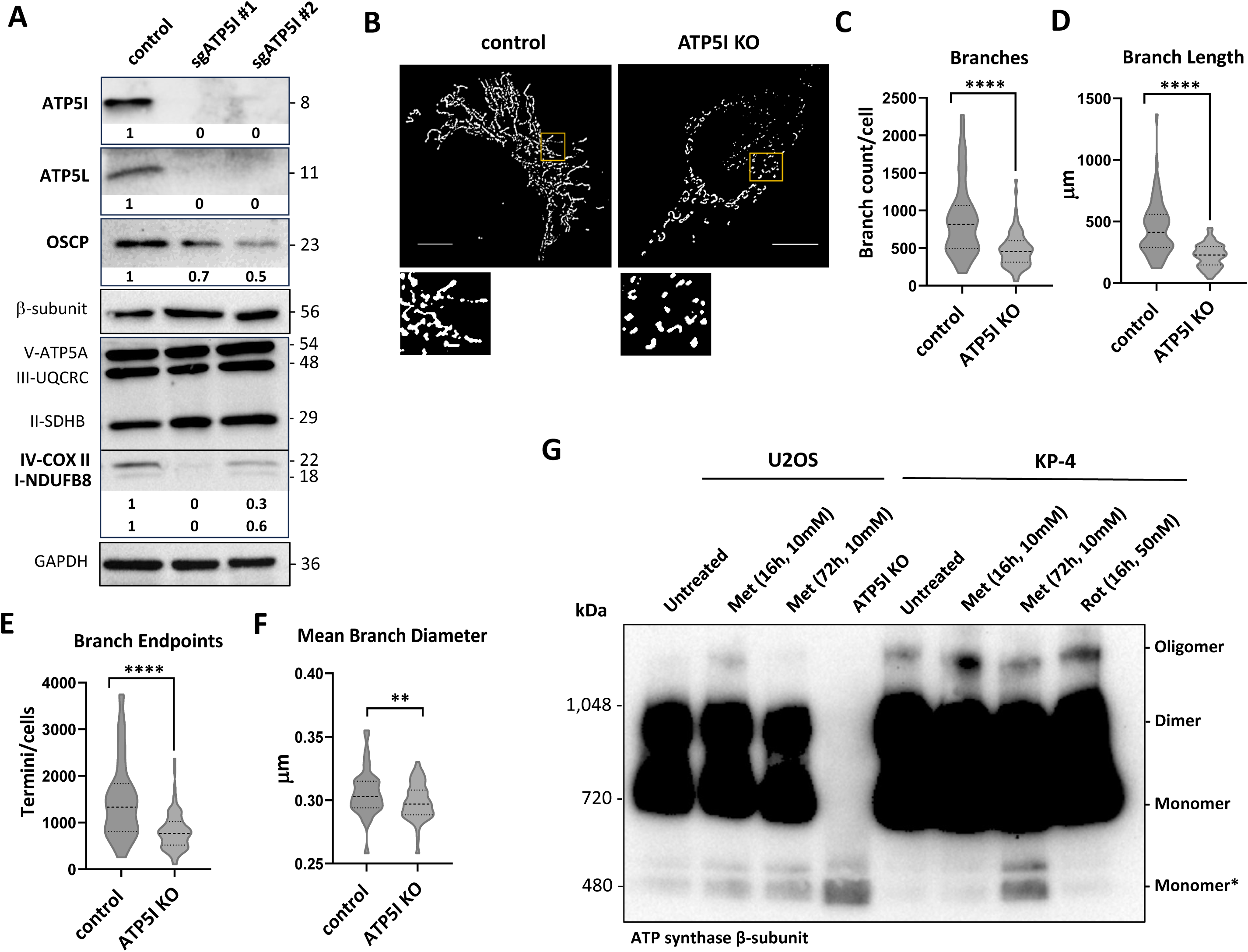
ATP5I knockout in pancreatic cancer cells alters the organization of the mitochondrial network. **(A)** Immunoblot for the indicated proteins in extracts from clones of KP-4 cells expressing a control small guide RNA against GFP (sgGFP: control 1 and control 2) or two different sgRNAs against ATP5I (sgATP5I #1 and sgATP5I #2). GAPDH antibody was used as loading control. **(B)** Representative threshold images of mitochondrial morphologies visualized by TOMM20 immunofluorescence (scale bar = 10 μm). A magnified inset (yellow box) is shown for each image to highlight mitochondrial structural details. All images were analyzed using the Mitochondria Analyzer plugin in Fiji (ImageJ). **(C–F)** Quantitative analysis of key mitochondrial parameters: (C) number of branches, (D) total branch length, (E) number of branch endpoints and (F) mean branch diameter. Data represent mean ± standard deviation from N = 3 independent clones, with 50–100 cells analyzed per clone. (ns) not significant, P < 0.01 (**), and P < 0.0001 (****) using an unpaired Student’s *t*-test. **(G)** Representative Blue Native-PAGE followed by western blotting using an antibody against the β-subunit of the F domain of ATP synthase in KP-4 or U2OS cells treated with vehicle, metformin (10 mM, 16 h or 3 days), or in ATP5I knockout cells (ATP5I KO). Monomer* indicates the assembly intermediates of the F F -ATP synthase known to accumulate after disabling ATP5I.

**Figure 2 – figure supplement 1.**
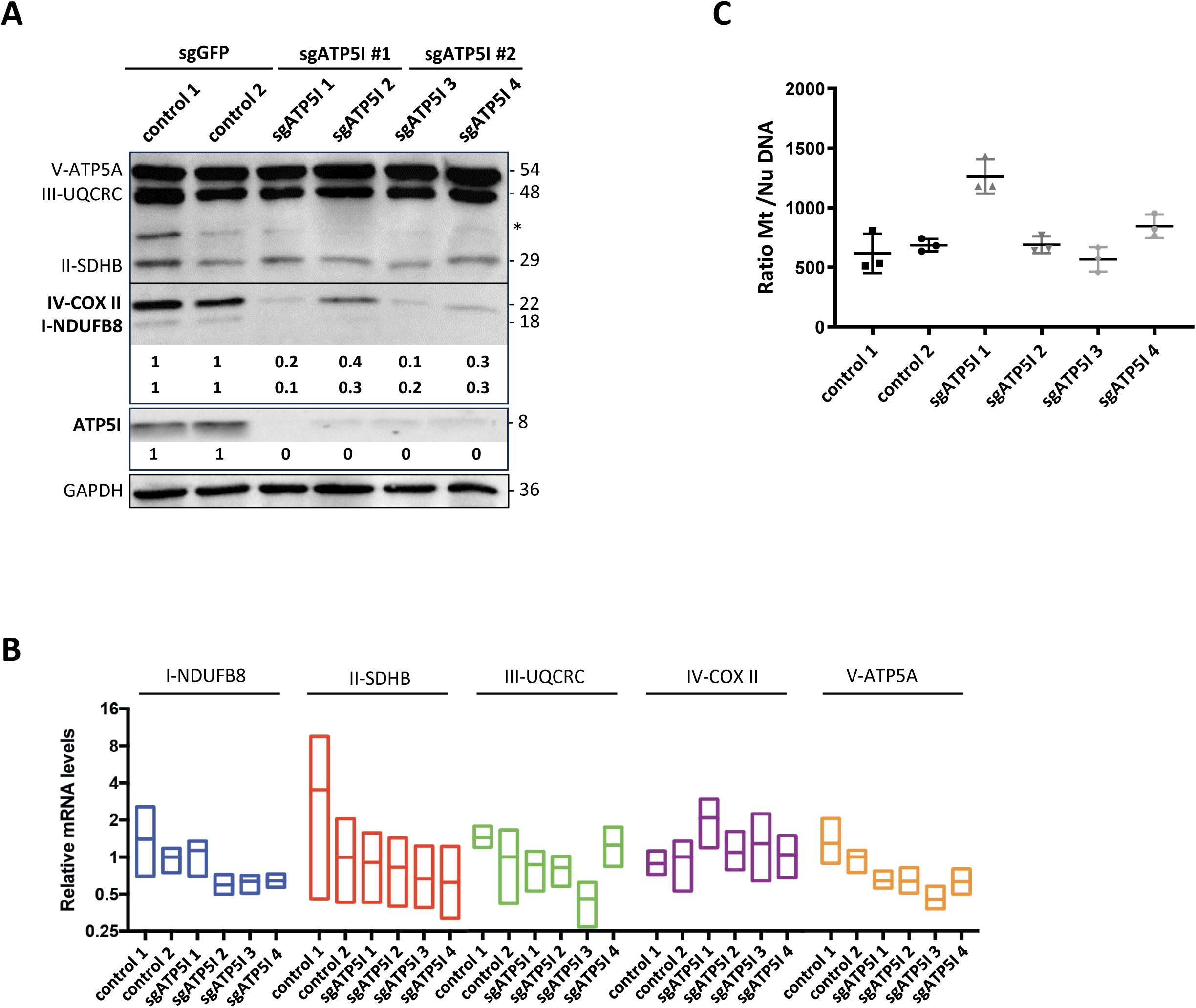
**(A)** Immunoblot for the indicated proteins in extracts from clones of KP-4 cells expressing a control small guide RNA against GFP (sgGFP: control 1 and control 2) or two clones for each of the two different guides targeting ATP5I (sgATP5I #1: sgATP5I 1 and sgATP5I 2, and sgATP5I #2: sgATP5I 3 and sgATP5I 4). * is a non-specific band that migrates like 40 KD MTCO1 from Complex IV. GAPDH was used as loading control. **(B)** Relative qPCR quantification of the mRNAs encoding proteins representative of the five OXPHOS complexes (in Figure 2A) in clones of KP-4 cells expressing a control small guide RNA against GFP (sgGFP: control 1 and control 2) or two clones for each of the two different guides targeting ATP5I (sgATP5I #1: sgATP5I 1 and sgATP5I 2, and sgATP5I #2: sgATP5I 3 and sgATP5I 4). Values represent the mean ± standard deviation of three biological replicates. **(C)** qPCR quantification of mitochondrial genomic DNA (Mt) over cellular nuclear genomic DNA (Nu) in cells as in (A). Values represent the mean ± standard deviation of N=3.

**Figure 2 – figure supplement 2.**
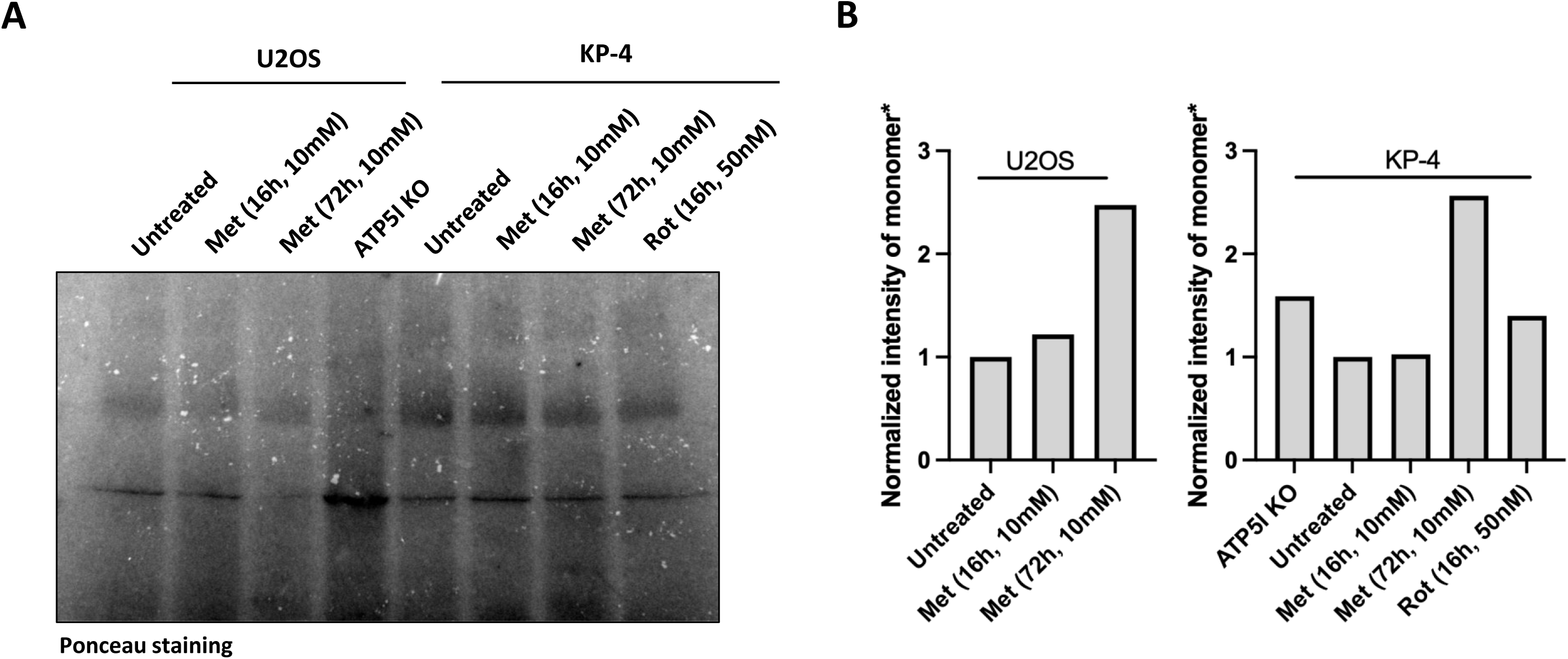
Quantification of immunoblot in Figure 2G. **(A)** The PVDF membrane was stained with Ponceau S to visualize total protein and subsequently imaged on Bio-Rad ChemiDoc XRS+ system. (**B)** Quantification. Using Bio-Rad Image Lab software, total band intensity was quantified for the band corresponding to the vestigial form of ATP synthase (monomer*) in figure 2G and normalized to the total band intensity of the corresponding lane in the Ponceau image. Values for each cell line were normalized to the “untreated” condition.

We then examined mitochondrial morphologies using fluorescence microscopy in ATP5I knockout (KO) cells, revealing a disruption in the mitochondrial network characterized by predominantly punctate mitochondria (Figure 2B-F), relative quantification showed that about ATP5I KO cells exhibited a punctate phenotype, characterized by less branches and branch end points and reduced diameter (Figure 2C-F). These findings confirm that ATP5I plays a crucial role in the organization of mitochondrial network in KP-4 cells. Treatment of the same cell line with medicinal biguanides resulted in similar alterations in mitochondrial network organization^37^ reinforcing the hypothesis that ATP5I may be a target of biguanides.

Finally, since ATP5I controls the dimerization and assembly of the F F -ATP synthase we investigated whether metformin treatment would affect this process. Supporting this model, BN-PAGE analysis of KP-4 cells treated with metformin (10 mM for 3 days) reveals a decrease in oligomeric forms and a corresponding accumulation of intermediate vestigial complexes of lower molecular weight than the monomeric enzyme as described in cells having CRISPR-mediated inactivation of ATP5I^53^ (Figure 2G and Figure 2 – figure supplement 2). Notably, this effect is not observed after short-term exposure (16 h), suggesting that the drug may inhibit the assembly of the F F -ATP synthase but do not disrupt already formed complexes. This increase in assembly intermediates of the F F -ATP synthase upon metformin treatment mimics the phenotype of ATP5I knockout cells where in addition the dimeric and monomeric forms of the enzyme are also scarce (Figure 2G). To broaden the significance of this finding, we reproduced it in U2OS cells (Figure 2G). Finally, rotenone, a complex I inhibitor, does not induce the formation of these vestigial complexes, indicating that the effect is not secondary to complex I alterations or reduced respiration induced by metformin.

### ATP5I knockout desensitizes pancreatic cancer cells to biguanides

Given the uncertain role of ATP5I in the cellular energy metabolism, we investigated the effects of ATP5I knockout (KO) on mitochondrial bioenergetics in KP-4 cells. The results reveal that ATP5I deletion significantly decreases the NAD^+^/NADH ratio (Figure 3A), most significantly affecting NAD^+^ concentration (Figure 3A – figure supplement 1A, 1B). Our results also indicate a decrease in the oxygen consumption rate (OCR) to extracellular acidification rate (ECAR) ratio (Figure 3B and Figure 3 – figure supplement 1C, 1D) suggesting a reduced respiration associated with a compensatory increase in glycolysis. This metabolic reorganization in ATP5I KO cells renders them up to 6 times more sensitive to glycolysis inhibition with 2-D-deoxyglucose (Figure 3 – figure supplement 1E), similar to control cells treated with 2.5 to 5 mM metformin (Figure 3 – figure supplement 1F). These findings confirm that both ATP5I deletion and metformin treatment disrupt respiration conferring a dependency on glycolysis^54^.

**Figure 3.**
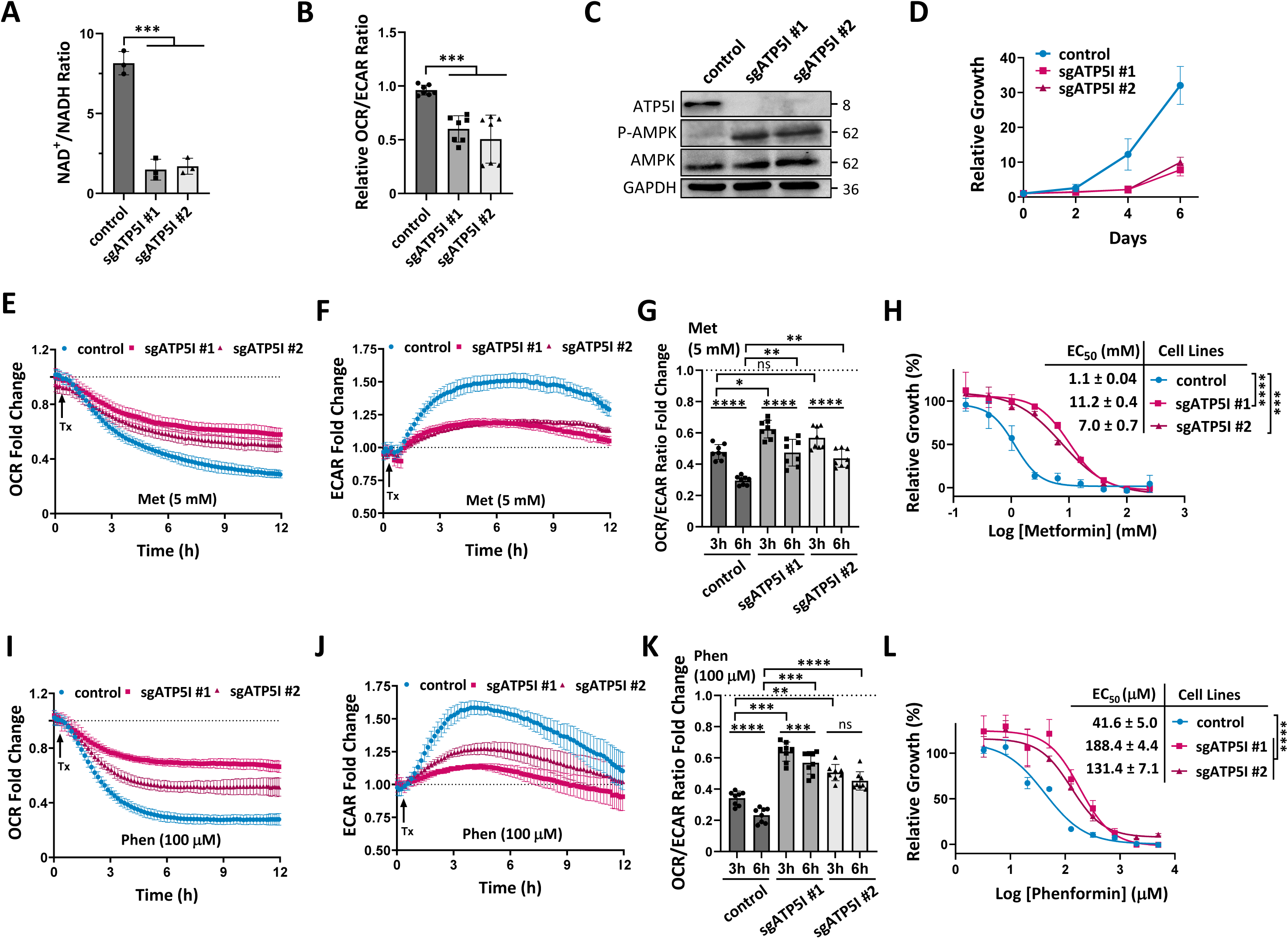
ATP5I knockout desensitizes pancreatic cancer cells to biguanides. **(A)** Quantification of NAD^+^/NADH ratio in KP-4 cells expressing a control small guide RNA against GFP (sgGFP) or a representative clone of two different guides targeting ATP5I (sgATP5I #1 or sgATP5I #2). Values represent the mean ± standard deviation of N=3. (***) P < 0.001 using an unpaired Student’s *t*-test. **(B)** Relative quantification of oxygen consumption rate (OCR) over extracellular acidification rate (ECAR) by Seahorse analysis in cells as in (A). Values represent the mean ± standard deviation of at least N=3. (***) P < 0.001 using a paired Student’s *t*-test. **(C)** Immunoblot for total and phosphorylated levels of AMPK (Thr172) protein in extracts from cells as in (A). ATP5I confirms loss of expression in KO and GAPDH was used as loading control. **(D)** Growth curves of cells as in (A) measuring the relative number of cells over 6 days. Media was changed every two days. **(E)** Representative kinetic curves of OCR in cells as in (A) treated with 5 mM of metformin (Met) relative to control treated cells (dashed line) using Seahorse. **(F)** Representative kinetic curves of ECAR in cells as in (A) treated with 5 mM metformin (Met) relative to control treated cells (dashed line) using Seahorse. **(G)** Quantification of OCR/ECAR ratio fold change at 3 and at 6 hours from kinetic curves (E-F). Values represent the mean ± standard deviation of N=3. (ns) not significative, (*) P < 0.05, (**) P < 0.01, (****) P < 0.0001 using a repeated measures (RM) one-way ANOVA with Sidak’s multiple comparison test. **(H)** Representative growth of cells as in (A) exposed to different concentrations of metformin (Met) for three days with corresponding EC_50_ values of metformin. Values represent the mean ± standard deviation of N=3. (***) P < 0.001 and (****) P < 0.0001 using an unpaired Student’s *t*-test. **(I)** Representative kinetic curves of OCR in cells as in (A) treated with 100 μM phenformin (Phen) relative to control treated cells (dashed line) using Seahorse. **(J)** Representative kinetic curves of ECAR in cells as in (A) treated with 100 μM of phenformin (Phen) relative to control treated cells (dashed line) using Seahorse. **(K)** Quantification of OCR/ECAR ratio fold change at 3 and at 6 hours from kinetic curves (I-J). Values represent the mean ± standard deviation of at least three biological replicates. ns: not significative, (**) P < 0.01, (***) P < 0.001, (****) P < 0.0001 using a RM one-way ANOVA with Sidak’s multiple comparison test. **(L)** Representative growth of cells as in (A) exposed to different concentrations of phenformin (Phen) for three days with corresponding EC_50_ values. Values represent the mean ± standard deviation of N=3. (****) P < 0.0001 using an unpaired Student’s *t*-test.

**Figure 3 – figure supplement 1.**
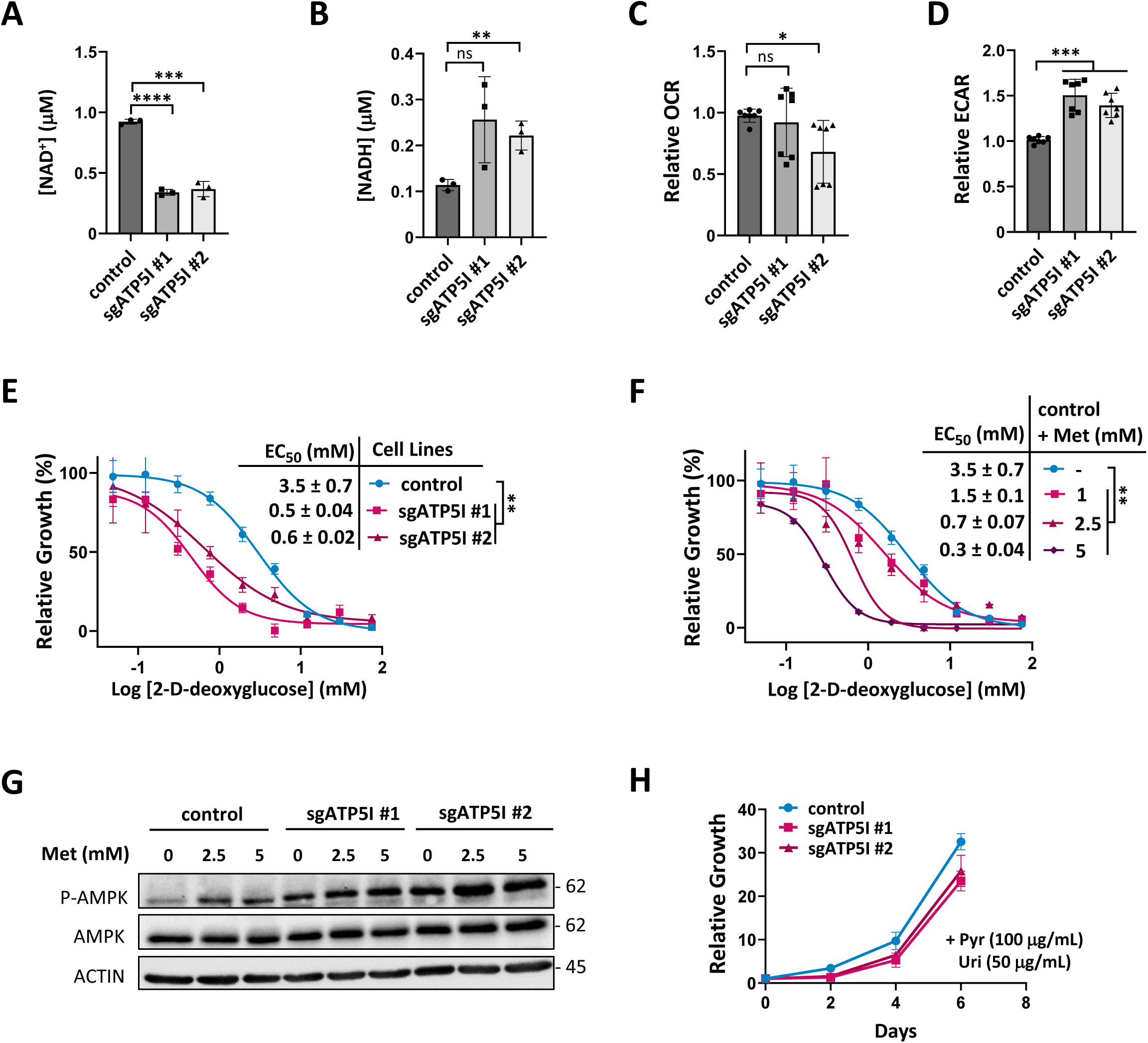
**(A)** Quantification of NAD^+^ concentration in KP-4 cells expressing a control small guide RNA against GFP (sgGFP) or a representative clone of two different guides targeting ATP5I (sgATP5I #1 or sgATP5I #2). Values represent the mean ± standard deviation of N=3. (***) P < 0.001, (****), P < 0.0001 using an unpaired Student’s *t*-test. **(B)** Quantification of NADH concentration in cells as in (A). Values represent the mean ± standard deviation of N=3. (ns) not significative, (**) P < 0.01 using an unpaired Student’s *t*-test. **(C)** Relative quantification of oxygen consumption rate (OCR) by Seahorse analysis in cells as in (A). Values represent the mean ± standard deviation of at least N=3. (ns) not significative, (*) P < 0.05 using a paired Student’s *t*-test. **(D)** Relative quantification of extracellular acidification rate (ECAR) by Seahorse analysis in cells as in (A). Values represent the mean ± standard deviation of at least N=3. (***) P < 0.001 using a paired Student’s *t*-test. **(E)** Representative cell growth of cells as in (A) treated with different concentrations of 2-D-deoxyglucose with corresponding EC_50_ values of 2-D-deoxyglucose treatment in cells as in (A). Values represent the mean ± standard deviation of N=3. (**) P < 0.01 using an unpaired Student’s *t*-test. **(F)** Representative cell viability curves with corresponding EC_50_ values of treatments of 2-D-deoxyglucose in combination without (-) and with different concentrations (1, 2.5 and 5 mM) of metformin (Met) in KP-4 cells expressing control sgGFP. Values represent the mean ± standard deviation of three biological replicates. (**) P < 0.01 using an unpaired Student’s *t*-test**. (G)** Total and phosphorylated levels of AMPK protein extracts from cells as in (A) treated with 2.5 mM or 5 mM of Met for 16 hours. β-ACTIN was used for loading control. **(H)** Growth curves of cells as in (A) supplemented with 100 μg/mL sodium pyruvate (Pyr) and 50 μg/mL uridine (Uri) by measuring the percentage of confluency over 6 days. Media was changed every two days.

Moreover, the inhibition of the mitochondrial respiratory chain observed in ATP5I KO cells induces AMPK activation (Figure 3C), similarly to control cells treated with metformin (Figure 3 – figure supplement 1G). However, the increased phosphorylation state of AMPK in ATP5I KO cells treated with metformin at these concentrations suggests that metformin can still activate AMPK via interactions with other targets such as PEN2^31^.

Energy stress resulting from ATP5I KO in KP-4 cells leads to a noticeable slowdown in cell growth (Figure 3D), although this effect can be reversed by supplementing the cell culture medium with pyruvate and uridine (Figure 3 – figure supplement 1H). These findings suggest that while ATP5I is not essential for growth in KP-4 cells, its deletion subtly affects energy metabolism, akin to the effects seen in KP-4 cells treated with medicinal biguanides^37^.

Seahorse kinetic monitoring of control and ATP5I KO cells treated with 5 mM metformin reveals, as expected, that metformin causes a decrease in oxygen consumption (Figure 3E) and an increase in glycolysis (Figure 3F) over time. However, these effects are more pronounced in control cells than what is observed in ATP5I cells, particularly in terms of glycolysis activation. The OCR/ECAR ratio also decreases more in control cells upon metformin treatment than in ATP5I KO cells (Figure 3G), suggesting reduced sensitivity to metformin in the KO cells. This resistance is also evident in the relative growth of cells under increasing doses of metformin. Indeed, ATP5I KO cells exhibit EC_50_ values up to 9 times higher than those of control cells (Figure 3H). Overall, ATP5I KO phenocopies metformin activity and blunts the additional effect of metformin suggesting that metformin acts partially through ATP5I.

Similar trends where obtained, when using 100 μM phenformin in seahorse kinetic with more pronounced changes in both respiratory activity decline (Figure 3I) and glycolysis activation (Figure 3J). These differences appear earlier but over a shorter time compared to metformin, possibly due to phenformin’s pharmacokinetic profile^55^. Again, phenformin affects both control and ATP5I KO cells but it does so less efficiently in ATP5I KO cells. The difference in OCR/ECAR ratio between control cells and the two ATP5I KO cell lines at 3 hours is significantly less impacted (Figure 3K), indicating lesser sensitivity of ATP5I KO cells to phenformin. Resistance to phenformin is also evident in cell growth, with ATP5I KO cells showing EC_50_ up to 4 times higher than those of control cells (Figure 3L). These results suggest that ATP5I mediates the metabolic and antiproliferative effects of biguanides in KP-4 cells.

### Re-expression of ATP5I normalizes mitochondrial morphology, metabolic profile and resensitizes ATP5I knockout pancreatic cancer cells to biguanides

To validate our previous hypotheses regarding the mechanism of action of metformin, we reintroduced wild-type ATP5I in ATP5I KO KP-4 clones. Immunoblot analysis results confirmed that exogenous ATP5I could be re-expressed in ATP5I KO cells, albeit at slightly reduced levels compared to control cells. Reintroducing ATP5I also restored the protein levels of ATP5L, OSCP, NDUFB8 and COX II in KP-4 ATP5I KO cells (Figure 4A). Furthermore, exogenous ATP5I colocalized with TOMM20, indicating its successful mitochondrial localization and led to a reorganization of the mitochondrial networks (Figure 4 – figure supplement 1). Indeed, quantitative analysis showed that ATP5I KO cells with exoATP5I exhibited an intermediate fragmented phenotype between punctate and filamentous forms (Figure 4B-F).

**Figure 4.**
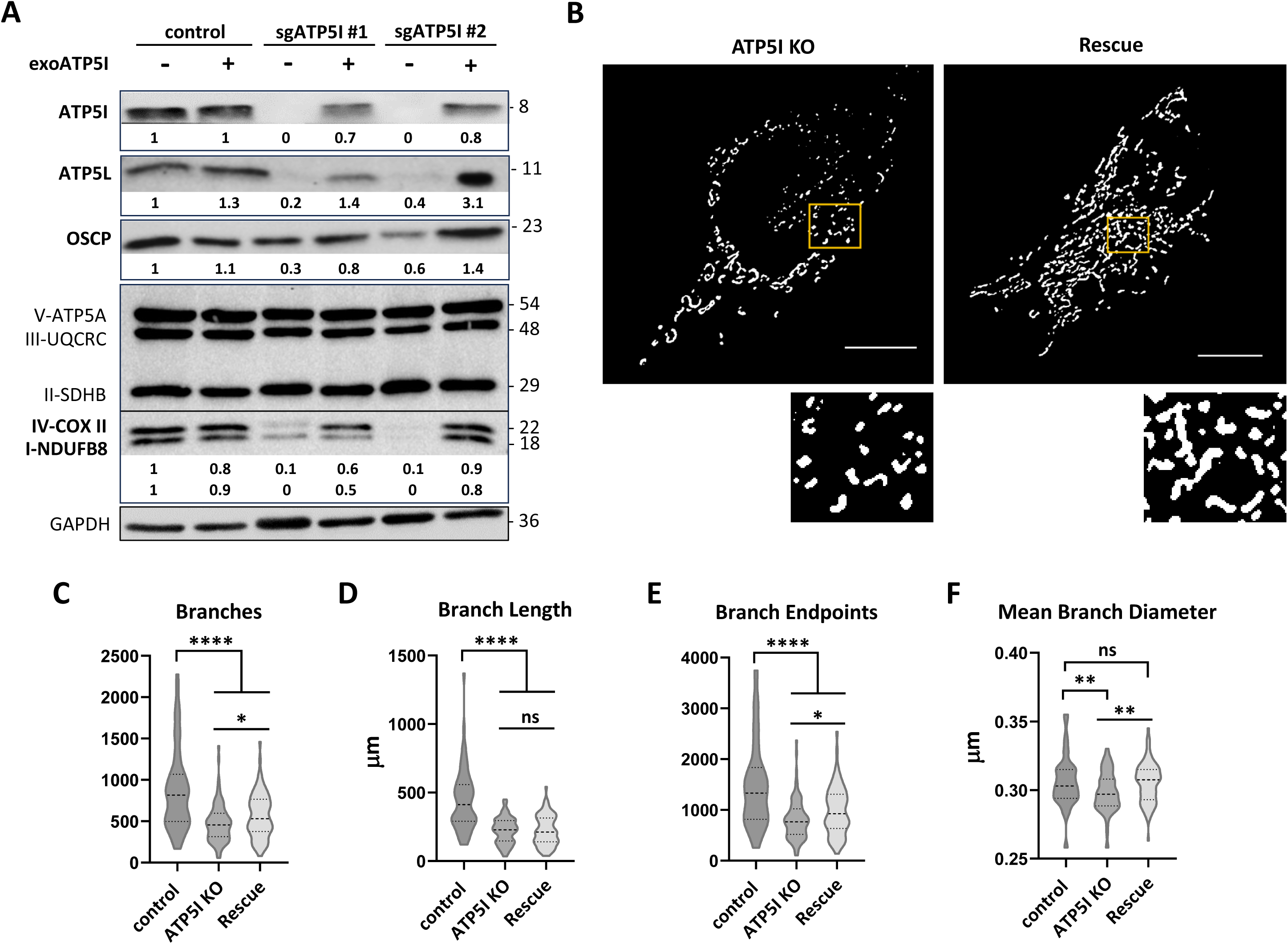
Exogenous ATP5I enable the reorganization of mitochondrial network in ATP5I knockout pancreatic cancer cells. **(A)** A representative immunoblots for the indicated proteins in KP-4 cells expressing exogenous ATP5I (exoATP5I: +) in control cells expressing a small guide RNA against GFP (sgGFP) or in ATP5I KO cells (clones of two different small guide RNAs: sgATP5I #1 sgATP5I #2) compared with the same cell lines without expression of exogenous ATP5I (-). GAPDH antibody was used as loading control. **(B)** Representative threshold images of mitochondrial morphologies visualized by TOMM20 immunofluorescence (scale bar = 10 μm) of ATP5I KO cells and their derivative re-expressing ATP5I. A magnified inset (yellow box) is shown for each image to highlight mitochondrial structural details. All images were analyzed using the Mitochondria Analyzer plugin in Fiji (ImageJ). **(C–F)** Quantitative analysis of key mitochondrial parameters: (C) number of branches, (D) total branch length, (E) number of branch endpoints and (F) mean branch diameter. Data represent mean ± standard deviation from N = 3 independent clones, with 50–100 cells analyzed per clone. (ns) not significant, P < 0.01 (**), and P < 0.0001 (****) using an unpaired Student’s *t*-test.

**Figure 4 – figure supplement 1.**
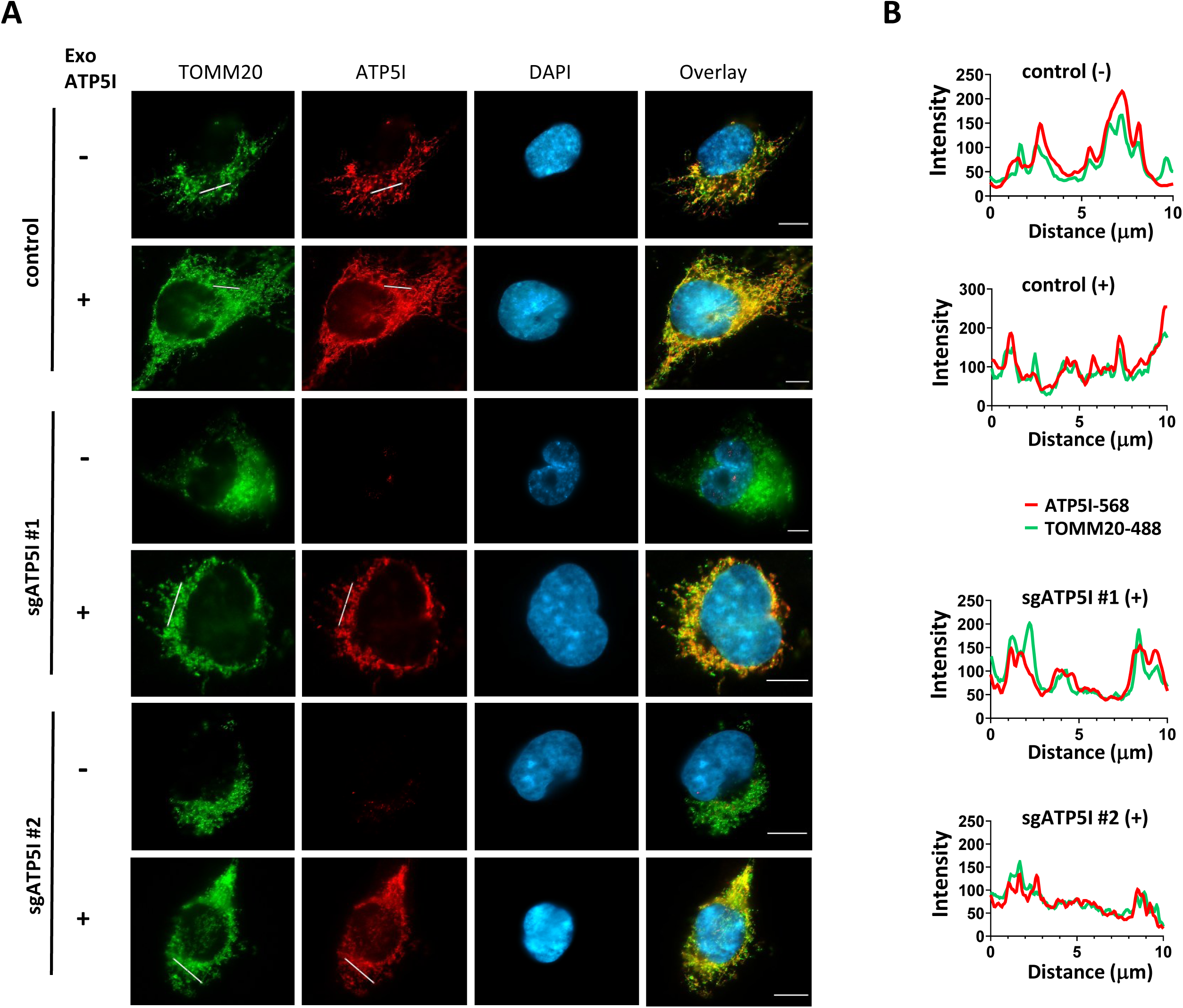
**(A)** Representative images of mitochondrial localization of ATP5I in KP-4 cells expressing exogenous ATP5I (exoATP5I: +) in control (clones with sgGFP) or a representative clone of each of two small guide RNA against ATP5I (sgATP5I #1 or sgATP5I #2) compared with the same cell lines without expression of exogenous ATP5I (-). Mitochondrial localization was analyzed by co-immunofluorescence using ATP5I and TOMM20 antibodies, scale bar= 10 μm. DAPI was used as a DNA counter stain. **(B)** Colocalization between TOMM20 (TOMM20-568) and ATP5I (ATP5I-488) signals was analyzed with job plot intensity profile.

Similarly, ATP5I KO cells expressing exogenous ATP5I showed increased NAD^+^/NADH ratios (Figure 5A) by restoring NAD^+^ concentration (Figure 5 – figure supplement 1A, 1B), increased OCR/ECAR ratio (Figure 5B), enhancing respiration and reducing compensatory glycolysis (Figure 5 – figure supplement 1C, 1D) to levels comparable to control cells. This metabolic reorganisation made these cells up to 3 times less sensitive to 2-D-deoxyglucose (Figure 5 – figure supplement 1E) compared to ATP5I KO cells, thereby alleviating energy stress as seen with less phosphorylated AMPK (Figure 5C), a tendency to increasing ATP levels (Figure 5D) and enabling increased cell growth (Figure 5E). Of note, ATP levels are not significantly different between the KO and control cells perhaps because an efficient compensatory glycolysis. Restoring the ATP5I expression is also correlated with increased sensitivity to the metabolic effects induced by metformin and phenformin on mitochondrial respiration and glycolysis and decreased OCR/ECAR ratios (Figure 5 – figure supplement 2). The rescue also enhanced the antiproliferative effects of metformin, with EC_50_ values up to 5-fold lower than those of ATP5I KO cells (Figure 5F), as well as those of phenformin, with EC_50_ values up to 3-fold lower than those of ATP5I KO cells (Figure 5G).

**Figure 5.**
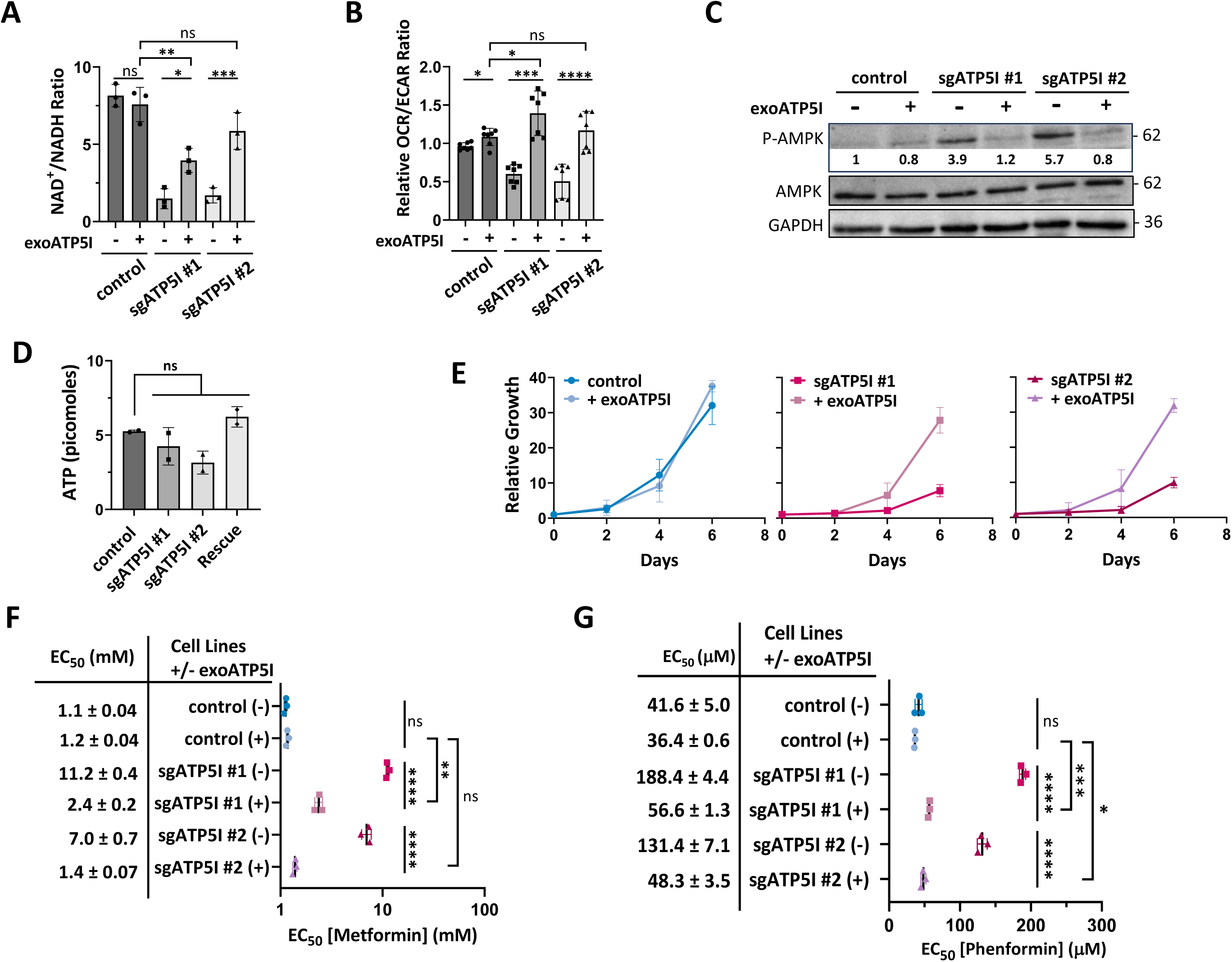
Re-expression of ATP5I rescues metabolic profile and resensitizes ATP5I knockout pancreatic cancer cells to biguanides. **(A)** Quantification of NAD^+^/NADH ratio in KP-4 cells expressing exogenous ATP5I (exoATP5I: +) in control sgGFP or a representative clone of two different small guide RNAs (sgATP5I #1 and sgATP5I #2) compared with the same cell lines without expression of exogenous ATP5I (-). Values represent the mean ± standard deviation of three biological replicates. (ns) not significative, (*) P < 0.05, (**) P < 0.01, (***) P < 0.001 using an ordinary one-way ANOVA with Sidak’s multiple comparison test. **(B)** Relative quantification of oxygen consumption rate (OCR) over extracellular acidification rate (ECAR) by Seahorse analysis in cells as in (A). Values represent the mean ± standard deviation of at least three biological replicates. (ns) not significative, (*) P < 0.05, (***) P < 0.001, (****) P < 0.0001 using a repeated measures (RM) one-way ANOVA with Sidak’s multiple comparison test. **(C)** Immunoblot of total and phosphorylated levels of AMPK (Thr172) protein in extracts from cells as in (A). GAPDH was used as loading control. **(D)** Intracellular ATP levels measured in cell lines as in (A). Data are presented as mean ± standard deviation. N = 2. ns: not significant. **(E)** Growth curves of cells as in (A) by measuring the relative number of cells over 6 days. Media was changed every two days. **(F)** EC_50_ values of metformin (Met) treatments in cells as in (A). Values represent the mean ± standard deviation of N=3. (ns) not significative, (**) P < 0.01, (****) P < 0.0001 using an ordinary one-way ANOVA with Sidak’s multiple comparison test. **(G)** EC_50_ values of phenformin (Phen) treatment in cells as in (A). Values represent the mean ± standard deviation of three biological replicates. (ns) not significative, (*) P < 0.05, (***) P < 0.001, (****) P < 0.0001 using an ordinary one-way ANOVA with Sidak’s multiple comparison test.

**Figure 5 – figure supplement 1.**
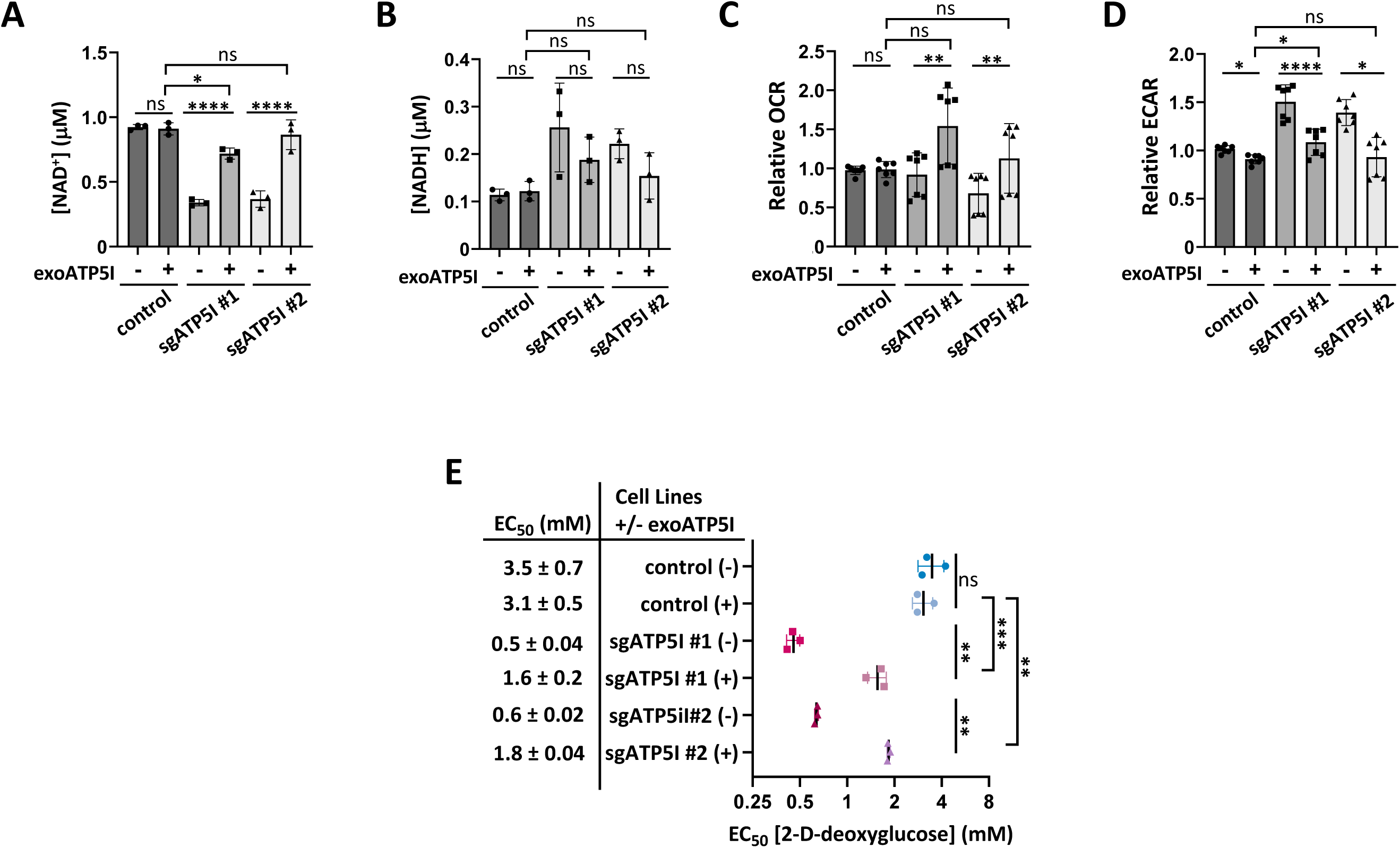
**(A)** Quantification of NAD^+^ concentration in KP-4 cells expressing exogenous ATP5I (exoATP5I: +) in control sgGFP clones or a representative clone of two different small guide RNAs against ATP5I (sgATP5I #1 and sgATP5I #2) compared with the same cell lines without expression of exogenous ATP5I (-). Values represent the mean ± standard deviation of three biological replicates. (ns) not significative, (*) P < 0.05, (****) P < 0.0001 using an ordinary one-way ANOVA with Sidak’s multiple comparison test. **(B)** Quantification of NADH concentration in cells as in (A). Values represent the mean ± standard deviation of three biological replicates. (ns) not significative using an ordinary one-way ANOVA with Sidak’s multiple comparison test. **(C)** Relative quantification of oxygen consumption rate (OCR) by Seahorse analysis in cells as in (A). Values represent the mean ± standard deviation of at least three biological replicates. (ns) not significative, (**) P < 0.01 using a repeated measures (RM) one-way ANOVA with Sidak’s multiple comparison test. **(D)** Relative extracellular acidification rate (ECAR) by Seahorse analysis in cells as in (A). Values represent the mean ± standard deviation of at least N=3. (ns) not significative, (*) P < 0.05, (****) P < 0.0001 using a RM one-way ANOVA with Sidak’s multiple comparison test. **(E)** EC_50_ values of 2-D-deoxyglucose treatment in cells as in (A). Values represent the mean ± standard deviation of N=3. (**) P < 0.01, (***) P < 0.001 using an ordinary one-way ANOVA with Sidak’s multiple comparison test.

**Figure 5 – figure supplement 2.**
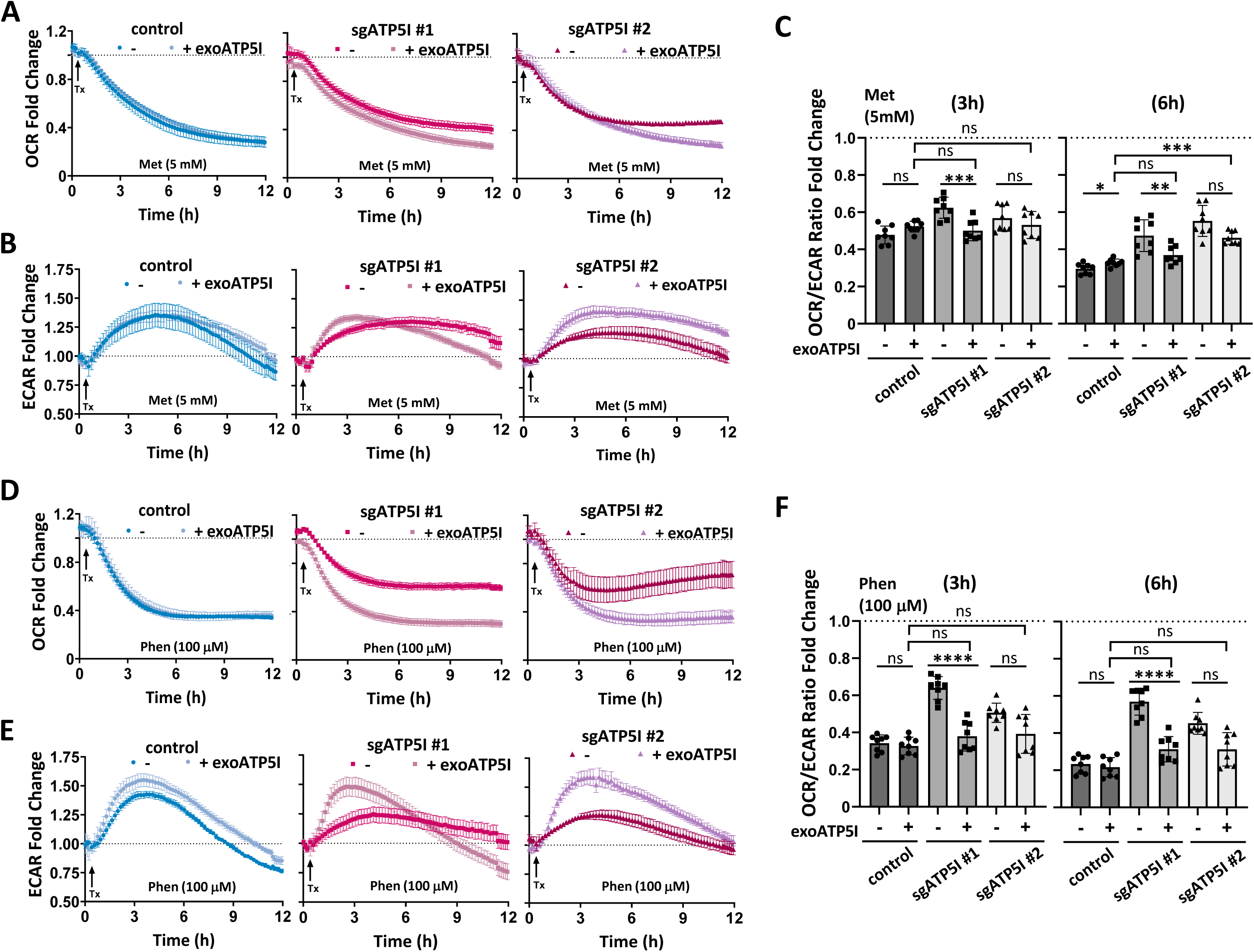
**(A)** Representative kinetic curves of oxygen consumption rate (OCR) fold change in KP-4 cells expressing exogenous ATP5I (exoATP5I: +) in control sgGFP clones or a representative clone of each of two small guide RNA against ATP5I (sgATP5I #1 and sgATP5I #2) compared with the same cell lines without expression of exogenous ATP5I (-) treated with 5 mM metformin (Met) using Seahorse. **(b)** Representative kinetic curves of extracellular acidification rate (ECAR) fold change in cells as in (A) treated with 5 mM Met using Seahorse. **(C)** Quantification of OCR/ECAR ratio fold change at 3 and at 6 hours from kinetic curves in (A) and (B). Values represent the mean ± standard deviation of at least three biological replicates. (ns) not significative, (*) P < 0.05, (**) P < 0.01, (***) P < 0.001 using a repeated measures (RM) one-way ANOVA with Sidak’s multiple comparison test. **(D)** Representative kinetic curves of OCR fold change in cells as in (A) but treated with 100 μM phenformin (Phen) using Seahorse. **(E)** Representative kinetic curves of ECAR fold change in cells as in (A) but treated with 100 μM Phen using Seahorse. **(F)** Quantification of OCR/ECAR ratio fold change at 3 and at 6 hours from kinetic curves (D) and (E). Values represent the mean ± standard deviation of at least three biological replicates. (ns) not significative, (****) P < 0.0001 using a RM one-way ANOVA with Sidak’s multiple comparison test.

In conclusion, these findings collectively demonstrate that the re-expression of ATP5I in ATP5I KO clones rescues mitochondrial morphology, metabolic profile and sensitivity to biguanides, which supports the concept that ATP5I mediates the metabolic and antiproliferative effects of these compounds in KP-4 cells.

### The Seahorse bioenergetic test reveals additional similarities between biguanides and ATP5I KO

To gain a deeper understanding of the individual contributions of respiratory complexes and the F F -ATP synthase in the cellular response to biguanides, we conducted the Seahorse Bioenergetic Mito Stress Test. This test measures baseline OCR and the response to specific inhibitors of mitochondrial complexes. Under basal conditions, treatment with biguanides at the indicated concentrations for 3 hours reduced the OCR in control cells, but less so in ATP5I KO cells, where OCR was already downregulated. Adding back ATP5I restored respiration and biguanide sensitivity to ATP5I KO cells (Figure 6A). Inhibition of the F F -ATP synthase with oligomycin reduced the OCR in control cells, but less so in biguanide treated cells or in ATP5I KO cells (Figure 6A). Following treatment with the uncoupler FCCP, respiration increases in biguanide-treated cells, and to a lesser extent in ATP5I KO cells (Figure 6A). Adding rotenone and antimycin A totally blocks this respiration confirming that FCCP requires active complexes I-IV to induce maximal respiration. Since defects in complex-I and other respiratory complexes are characterized by a lack of response to uncouplers^56,57^, our results are consistent with either a direct but partial inhibitory activity of biguanides on the complexes or targeting the F F -ATP synthase by biguanides and a subsequent disorganization of the electron transport chain and cristae (indirect mechanism). These effects are even more pronounced in ATP5I KO cells, where F F -ATP synthase oligomerization is completely inhibited (Figure 2G).

**Figure 6.**
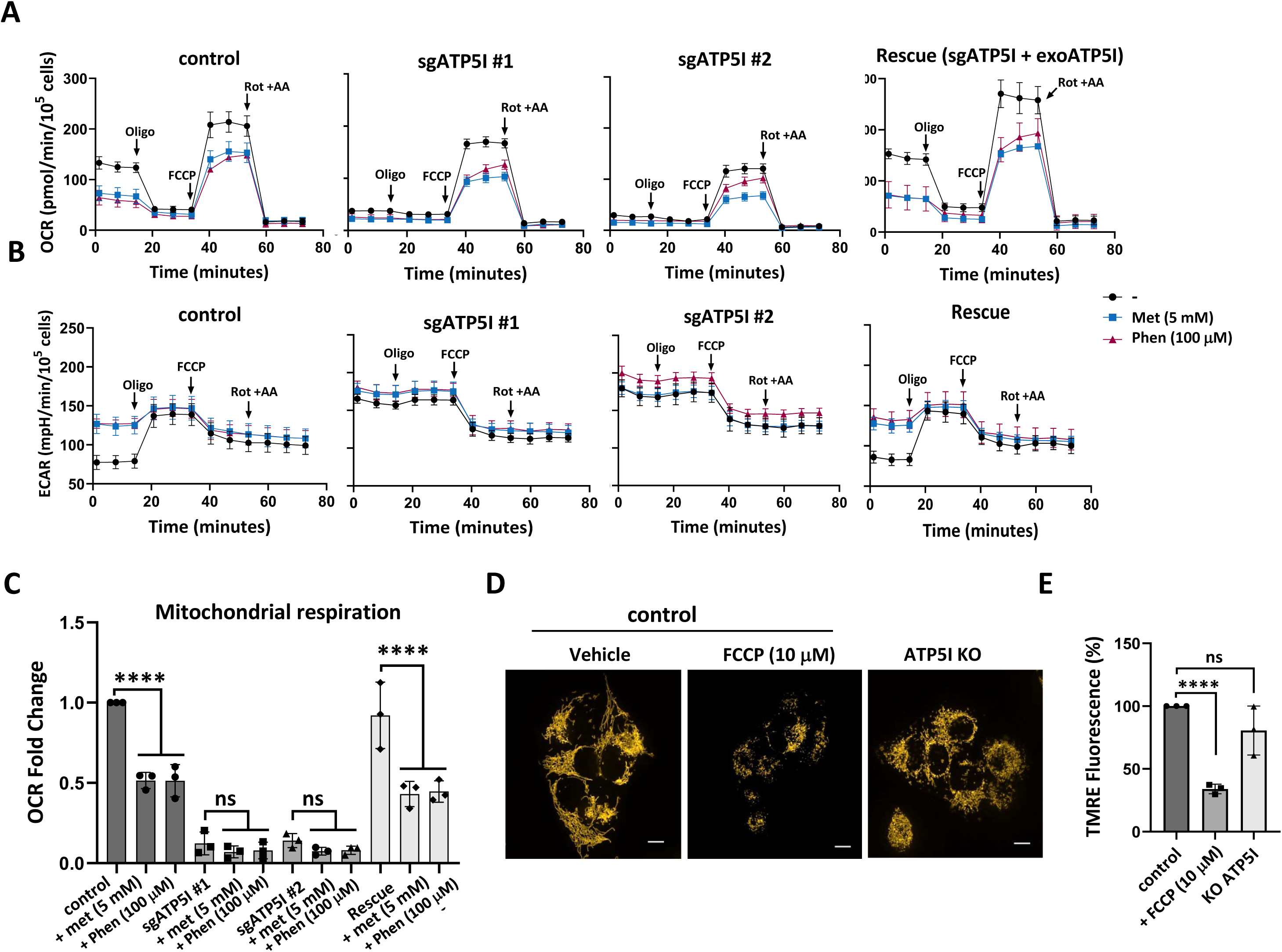
ATP5I Deletion Mimics Biguanide-Induced Bioenergetic Remodeling. **(A)** Representative oxygen consumption rate (OCR) profiles following sequential injection of oligomycin (Oligo), FCCP, and rotenone/antimycin A (Rot/AA) in control, ATP5I knockout (sgATP5I #1 and #2), and Rescue (sgATP5I + exoATP5I) cells treated with vehicle, 5 mM metformin (Met), or 100 µM phenformin (Phen). **(B)** Representative extracellular acidification rate (ECAR) profiles under the same conditions as in (A). **(C)** Quantification of basal mitochondrial respiration corresponding to the conditions in (A), N=3. **(D)** Representative confocal images of cells stained with the membrane potential sensitive dye TMRE (100 nM, 30 min at 37 °C in complete DMEM without phenol red) under control conditions, following ATP5I knockout (ATP5I KO), or after depolarization with FCCP (10 µM, 30 min prior to TMRE incubation), scale bar = 10 µm. **(E)** Quantification of TMRE fluorescence intensity per cell. Data are expressed as mean ± standard deviation and normalized to control levels. Results are from three independent experiments performed on separate days (n = 431 cells for control; n = 371 for FCCP; n = 536 for ATP5I KO). (ns) not significative and (***) P < 0.001, (****) P < 0.0001 using an unpaired Student’s *t*-test.

Analysis of ECAR reveals elevated basal glycolytic activity both in biguanide treated and in ATP5I KO cells (Figure 6B). Moreover, while oligomycin typically increases glycolysis in control cells by inhibiting F F -ATP synthase and promoting compensatory glycolytic ATP production, it has little to no effect in ATP5I-deficient cells or in biguanide-treated cells (Figure 6B). This suggests that biguanide treatment or ATP5I loss by impairing complex V function, render cells less responsive to oligomycin. Importantly, the reintroduction of ATP5I (rescue model) restored glycolytic activity to levels comparable to those of control cells (Figure 6B). Of note biguanides inhibited OCR by 50% in control cells but did not significantly reduced respiration in ATP5I KO cells (Figure 6C). Adding back ATP5I to KO cells restored the biguanide sensitivity of OCR (Figure 6C).

Finally, one could argue that biguanides have a lesser effect in ATP5I KO cells due to a disruption in membrane potential which is required for metformin to enter mitochondria. We thus used TMRE (tetramethylrhodamine ethyl ester) to measure the mitochondrial membrane potential in ATP5I KO cells and found it to be indistinguishable from that of control cells expressing wild-type ATP5I, although markedly higher than in cells treated with the uncoupler FCCP that reduced it by 65% (Figure 6D-E). These results align with the morphological changes detailed in Figure 2 and indicate that ATP5I deletion compromises mitochondrial structure without causing a profound loss of membrane polarization. Together, these findings indicate that while both FCCP treatment and ATP5I deletion disrupt mitochondrial morphology, only FCCP leads to a pronounced loss of membrane potential. These results support our proposition that biguanides require ATP5I to inhibit mitochondrial function.

### Chemogenomic screening of metformin reveals an imprint on F F -ATP synthase

Several candidate targets have been reported for biguanides and our results presented so far suggest a new one. Clues about drug mechanism of action can be obtained in impartial manner using genetic perturbation^58^. To obtain unbiased observation of biological processes affected by metformin, we performed a genome-wide pooled CRISPR/Cas9 KO screen in NALM-6 cells cultured in the presence of metformin at a concentration affecting growth (16 mM). Over the 8-day period of screening, cells had 1.47 population doublings whereas untreated controls had 7.5 (a 65-fold difference in growth). Based on the sgRNA frequency changes observed afterward between treated vs untreated samples, we generated enrichment/depletion scores for each gene using the CRANKS algorithm. Many gene KO were then predicted to potentiate (sgRNA depletion, KO creates sensitivity, negative CRANKS scores) or to suppress (sgRNA enrichment, KO creates resistance, positive CRANKS scores) metformin-induced growth inhibition (Figure 7A). Inactivation of the amino acid transporter SLC7A6 that plays a role in arginine export^59^ enhanced metformin activity, likely because this protein may also export metformin. Indeed, the end of the R group is very similar chemically to metformin. In addition, disabling the transcription factors SOX4, TCF4, SPI1 and the histone H3K79 methyltransferase DOT1L also enhanced metformin activity perhaps because they promote gene expression programs that compensate the mitochondrial dysfunction associated to metformin action. For example, SOX4 and SPI1 can increase glycolysis by promoting HK2 expression^60,61^ while TCF4 increases glycolysis by inducing the expression of the glucose transporter GLUT3^62^. Moreover, metformin can increase DOT1L mediated H3K79 methylation promoting the expression of SIRT3, a mitochondrial deacetylase that stimulates mitochondrial health and biogenesis^63^. On the other hand, many genes were found to be required for metformin-mediated growth inhibition. Of note, DDX3X, an RNA helicase required for the translation of interferon response genes^64^, suggests a role for the interferon pathway in the antiproliferative activity of metformin. The actions of metformin were also suppressed by inactivation of the OMA1-DELE1-HRI pathway. OMA1, is a protease localized to the inner mitochondrial membrane activated by mitochondrial stress. DELE1 is a target of OMA1 that interacts with and activates EIF2AKI (also known as heme-regulated inhibitor of HRI). HRI catalyzes eIF2a phosphorylation which in turns activates the translation of ATF4^65^ which is also stimulated by DDX3X^66^. ATF4 together with ATF3 regulates the expression of prop-apoptotic genes in response to both ER and mitochondrial stress^65,67^. Finally, inactivation of VDAC2, BAK1, HCCS (which makes Cytochrome C), Cytochrome C (CYCS) and APAF1 also conferred resistance to metformin, suggesting that metformin induces the classical mitochondria-related intrinsic apoptosis in treated cells. We compared the pattern of genetic interactions of metformin with that of rotenone, a complex I inhibitor (Figure 7B) and oligomycin A, a complex V inhibitor (Figure 7C). Interestingly, there were more similarities between oligomycin A and metformin (16 genes in common) than between rotenone and metformin (4 genes in common). Notably, inactivation of the mitochondrial apoptotic pathway through OMA1-DELE1-HRI as well as glycolytic pathway through SOX4 and TCF4 significantly affected both oligomycin A and metformin but not rotenone. On the other hand, DDX3X was required for the actions of all three drugs. Taken together, this genetic experiment shows that the OMA1-DEL1-HRI pathway mediates the antiproliferative activity of biguanides or the F_1_F_o_-ATP synthase inhibitor oligomycin (Figure 7D) while a compensatory glycolysis protects the cells.

**Figure 7.**
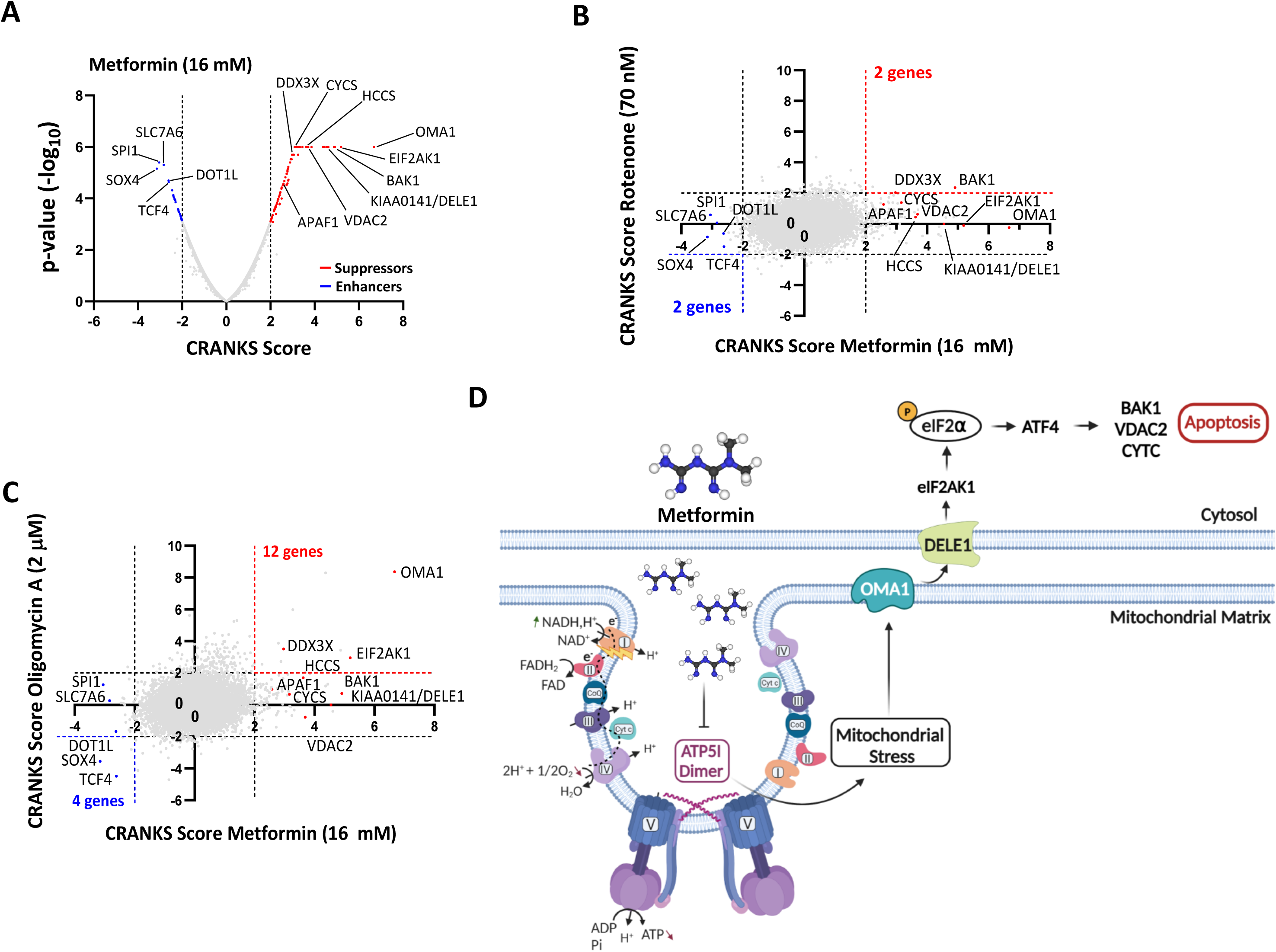
Chemogenomic screening of metformin reveals an imprint on F_1_F_o_-ATP synthase. **A)** Results of the pooled genome-wide CRISPR/Cas9 KO screen made in NALM-6 cells treated with 16 mM metformin or control. Data are represented as a Volcano plot of gene enrichment/depletion scores vs p-values from using the CRANKS algorithm. Some genes of interest are labeled. Enhancers of metformin growth inhibition with negative CRANKS scores below 2.5 (dashed line) are labeled blue, while suppressors with positive CRANKS scores above 2.5 (dashed line) are labeled red. **B-C)** Pairwise comparison of gene CRANKS scores obtained from screening metformin 16 mM in NALM-6 cells against that from screening either 70 nM rotenone or **(C)** 2 μM oligomycin A. D) Model for metformin action triggering the OMA1-DELE1-HRI pathway.

## Discussion

Using a biologically active, biotin-functionalized biguanide (BFB), we identified and characterized the ATP synthase subunit e (ATP5I; Complex V) as a candidate mitochondrial target underlying the antineoplastic activity of medicinal biguanides. Biguanides bound purified ATP5I with low-micromolar affinity, and in cells they disrupted F F -ATP synthase oligomerization, accompanied by the appearance of lower–molecular weight vestigial intermediates. Consistent with Complex V involvement, biguanide-treated cells were able to reinitiate respiration upon uncoupling, and this recovered respiration was abolished by Complex I and III inhibitors, supporting a model in which biguanides impair ATP synthase organization/function while leaving upstream electron transport capacity intact.

Notably, several studies have argued that Complex I inhibition accounts for the mitochondrial effects of biguanides. In one report, uncouplers restored respiration in metformin-treated cells—an observation we also make here—and the authors proposed that uncoupling prevents metformin entry into mitochondria by collapsing the membrane potential required for its uptake^24^. However, we observed that uncoupling also restored respiration in phenformin-treated cells, even though phenformin does not require membrane potential to accumulate within mitochondria. Another study similarly found that uncouplers restored respiration in metformin-treated pancreatic cancer cells grown in adherent conditions, but not when the same cells were grown as spheroids, where the uncoupler failed to stimulate respiration, as happens in cells treated with rotenone^68^. These context-dependent findings are compatible with the idea that electron transport is influenced not only by the intrinsic activity of respiratory complexes, but also by their organization within the inner mitochondrial membrane and cristae—an architecture that depends on F F -ATP synthase oligomerization.

We noticed that biguanides have an immediate effect on mitochondrial respiration, but we needed to expose cells for longer times (72h in KP-4 cells) to observe an accumulation of vestigial intermediates of the F F -ATPase. Although the accumulation of these intermediates is consistent with biguanides affecting ATP5I functions, the explanation for the inhibition of respiration is still unknown. It is plausible that biguanides perturb functions of the peripheral stalk of the F F -ATP synthase, a structurally essential but still poorly characterized module with potentially broad effects on mitochondrial electron transport and respiratory activity. In addition, ATP5I has been reported to interact with and modulate proteins involved in Complex I import and assembly, including TIMMDC1^69,70^, as well as TMEM70 and TMEM242^71^. It is therefore plausible that biguanides could impair Complex I indirectly by disrupting ATP5I-dependent organization and/or the function of ATP5I partners. Consistent with this model, both biguanide treatment and ATP5I knockout lead to a reduced NAD /NADH ratio, indicating functional impairment of mitochondrial complex I.

Inhibition of respiration can disrupt the mitochondrial membrane potential, which is essential—among other functions—for the accumulation of metformin within mitochondria. Hence, a loss of membrane potential could explain the resistance to metformin in ATP5I KO cells. However, inhibition of respiration can trigger the F F -ATPase to operate in reverse, utilizing glycolytic ATP to sustain the membrane potential and thereby allowing continued mitochondrial uptake of metformin^24^. We confirmed this interpretation since both biguanide-treated cells and ATP5I KO cells exhibit compensatory glycolysis to maintain ATP levels close to normal and the mitochondrial membrane potential we measured in ATP5I KO cells was similar to control cells. In addition, phenformin, does not require cationic transporters and the membrane potential to inhibit mitochondrial functions^72^ and our ATP5I KO cells were also resistant to this drug (Figure 5G).

Similar to medicinal biguanides effects on KP-4 cells^37^, ATP5I depletion leads to mitochondrial structural changes such as mitochondrial fragmentation. Additionally, ATP5I deletion disrupts energy metabolism, decreasing complex I mediated NADH reoxidation and respiration, promoting glycolysis and AMPK activation, and impairing cell growth. Furthermore, ATP5I KO cells exhibit resistance to metformin and phenformin growth effects, indicating its pivotal role in mediating their metabolic and antiproliferative effects. Reintroducing wild-type ATP5I in ATP5I KO cells partially restored their sensitivity to biguanides, consistent with the proposed role of ATP5I being a biguanide target. Of note, the phenotypes observed in the absence of subunit g (ATP5L)^73^, the direct partner of ATP5I, and subunit f (ATP5K)^74^ in HeLa cells are similar to those of ATP5I mutants and metformin treated cells. Further work is needed to determine how metformin affects the interactions between ATP5I, ATP5L and ATP5K. Together these three proteins are located at the base of F_1_F_o_-ATP synthase peripheral stalk which interact with OSCP (oligomycin sensitivity conferring protein) located in the F_1_ portion of the enzyme ^45^. ATP5I’s movement within F_1_F_o_-ATP synthase^45^ may regulate and transfer information between the F_1_ and the F_o_ rotating complex ^42,69–71^. These structural connections could underpin the results reported here that metformin and oligomycin share a common genetic signature in a CRISPR screening in NALM-6 cells.

Our findings, combined with recent published results, strengthen the correlation between mitochondrial ultrastructure and respiratory activity^75^, suggesting that medicinal biguanides partially affect mitochondrial respiratory chain activity by indirectly inhibiting complex I of OXPHOS through ATP5I binding, although the exact complete molecular mechanism remains unclear. Interestingly, in our screening, metformin’s effects on cell growth were inhibited by the knockout of the mitochondrial protease OMA1. The expression of the complex I protein NDUFB8 was reduced in our ATP5I KO cells. Since NDUFB8 is a target of the mitochondrial protease ClpP, whose activation can inhibit pancreatic adenocarcinoma growth^76^, these results suggest the intriguing possibility that biguanides may exert anticancer effects by activating mitochondrial proteases.

The upregulation of ATP5I observed in hypoxia^77^, DNA damage^78^, liver cancer^79^, and low-fat diet responses^80,81^ underscores a regulatory role still poorly understood. In another study, RNA-seq data form A549 cells also suggest ATP5I levels influence metformin sensitivity^82^, highlighting its potential clinical relevance. Identifying ATP5I as a potential antineoplastic target for medicinal biguanides provides insights into the mechanisms underlying their anticancer effects and opens new avenues for targeting mitochondrial metabolism. These findings also underscore the need for further exploration of ATP5I’s functions within F F -ATP synthase and its connections with other OXPHOS complexes, potentially leading to novel therapeutic strategies for cancer treatment.

## Acknowledgements

We wish to express our deepest gratitude to Professor Omichinski, for his expert guidance and unwavering support in the purification of proteins, and also to Professor Masson and Affinité Instrument for their generosity in providing access to the SPR system. SPG is supported by Grants 840633, 1054571 from The Cancer Research Society. ARS by Cancer Research Society (CRS) and Charlotte Légaré Memorial Fund 935858 and NSERC RGPIN-2021-03128. GF by Grants form the Terry Fox Research Institute and The Cancer Research Society and the CIBC chair for breast cancer research.

## Author contributions

The manuscript was drafted by G.L., with all authors approving the final version. The project was initially launched by M.C.R. and F.M., who performed the first pull-down experiments (Fig. 1B and 1F). V.B. generated the stable ATP5I knockout cells and conducted all qPCR analyses (Fig. 2 – figure supplement 1B, 1C). E.L. carried out the Seahorse experiments (Fig. 3B, 3E, 3F, 3G, 3I, 3J, 3K, 5B; Fig. 3 – figure supplement 1C, 1D; and Fig. 5 – figure supplement 2). A.M.D. and M.N performed the BN-PAGE analyses (Fig. 2G and Figure 2 – supplementary figure 2). T.B. and M.K. performed the chemogenomic CRISPR/Cas9 knockout screen (Fig. 7). G.L. completed the remaining experimental work. A.R.S., G.F., and S.P.G. supervised the project and contributed to the finalization of the manuscript. All authors declare no competing interests.

## Materials and methods

### Drugs

Metformin hydrochloride was purchased from Combi-Blocks (#ST-9194; San Diego, CA, USA), phenformin hydrochloride was purchased from Sigma Aldrich (#P7045; Oakville, On, Canada), 2-D-deoxyglucose was purchased from Bioshop (#DXG498.5; Burlington, On, Canada) and D-biotin was purchased from Invitrogen/Thermo Fisher Scientific (#B1595; Waltham, MA, USA).

### Synthesis

All chemicals were purchased from Sigma Aldrich, Oakwood Chemicals (Estill, SC, USA) and Combi-Blocks in their highest purity and were used without further purification. Deuterated dimethylsulfoxide (DMSO-*d*6), deuterated methanol (MeOD) and deuterium oxide (D2O) were purchased from CDN Isotopes (Pointe-Claire, Qc, Canada). NMR spectra were recorded on Bruker avance 400 and Bruker avance 500 spectrometers. Coupling constants (J) were reported in hertz (Hz), chemical shifts were reported in parts per million (ppm, δ) and multiplicities were reported as singlet (s), doublet (d), triplet (t) and multiplet (m). 1H and 13C chemical shifts are relative to the solvents: δH 2.50 δC 39.5 for DMSO-*d*6, δH 3.35, 4.78 and δC (49.3) for MeOD and δH 4.65 for D2O. Final compounds were purified using a preparative liquid chromatography (prepLC-MS-Quadrupole, Waters) equipped with a C18 reverse phase column (2.1 x 100 mm, 3 mm, Atlantis) with HPLC-grade solvents (Sigma Aldrich). High-resolution mass spectra (HRMS) and LC-MS purity analysis were collected using LC-TOF with ESI ionisation source (Agilent Technologies, Santa Clara, CA, USA) by the regional mass spectrometry center of University of Montreal. After prep HPLC, all final compounds were subsequently freeze-dried and aliquots of the monochloride salt powder were carefully weighted. The detailed synthetic procedures and compound characterization are provided in the Supporting Information.

### Cell culture

Human pancreatic ductal carcinoma cell line KP-4 was as a kind gift from Dr. N. Bardeesy (Massachusetts General Hospital, Boston). Human embryonic kidney HEK-293T cells were obtained from Thermo Fisher Scientific. Phoenix ampho packaging cells was as a kind gift from Dr. S.W. Lowe (MSK, New York). NALM-6 cells were a gift from Stephen Elledge (Harvard, USA). U2OS cells were purchased from ATCC. Except for NALM-6 cells which were cultured in RPMI medium (#350-035; Wisent, all cells were cultured in Dulbecco’s modified Eagle medium (DMEM #319-015; Wisent, St-Jean-Batiste, Qc, Canada) supplemented with 10% fetal bovine serum (FBS, Wisent), 2 mmol/L L-glutamine (Wisent) and 1% penicillin/streptomycin sulfate (Wisent) in a humidified incubator at 37 °C with 5% CO_2_.

### Immunoblotting

For immunoblotting, proteins extracts were obtained from 6-well cell culture dishes at 80-90 % confluence. Cells were treated with the corresponding drugs or vehicle 16 hours after seeding. As previously described^83^, cells were washed twice with ice-cold phosphate-buffered saline solution (PBS), then the residual PBS was removed by aspiration. Cells were lysed in 250 μL of 2x Laemmli buffer (4% SDS, 20% glycerol, 120 mM Tris-HCl pH 6.8), recovered using a cell scraper and transferred into Eppendorf tubes for a mild sonication of 20s followed by heating for 5 min at 95 C. Protein extracts were cooled to room temperature (rt) and quantified by measuring absorbance at 280 nm using Nanodrop microvolume spectrophotometer (2000c, Thermo Fisher Scientific). Samples were then diluted to a final concentration (2 mg/mL) using a modified Laemmli Buffer (10% β-mercapthoethanol, 0.1% bromophenol, 2x Laemmli) and were kept at -20 C until use. A quantity of 25 to 40 μg of protein extracts were loaded into SDS-PAGE (12-15% for resolving gels, 4% for stacking gels) and run in Tris-Glycine SDS buffer (192 mM glycine, 0.1% SDS, 25 mM Tris-Base) at 90 V. SDS-PAGE were then transferred on a nitrocellulose membrane (0.45 µm, Bio-Rad, Mississauga, On, Canada) in Tris-Glycine buffer (96 mM glycine, 10 mM Tris-Base) for 1h30 at 120 V. Membranes were blocked in a modified Tris-buffered saline + 0.05% Tween-20 (TBST) solution with 5% skim milk for 1h at rt, washed with TBST 3x 5 min at rt and were then incubated with an appropriate diluted solution of primary antibody (in 0.1% BSA, 0.02% sodium azide, PBS pH 7.4) overnight (o/n) at 4°C. After primary antibody incubation, membranes were washed with TBST 3x 5 min and incubate with a diluted (1:3000 in 1% skim milk/TBST) secondary antibody coupled HRP for 1 hour at rt. After secondary antibody incubation, membranes were washed with TBST 3x 5 min. The signal was revealed using enhanced chemiluminescence reagent (ECL), (#RPN2106, GE Healthcare Life Sciences, Chicago, IL, USA) and images were acquired by exposition with autoradiographic films or using ChemiDoc Imaging Sytems (XRS+; Bio-Rad). FroggaBio BLUelf Prestained Protein ladder (FroggaBio, Concord, Om, Canada) was used to estimate protein molecular weight. Of note, most membranes were cut into pieces and were incubated with several antibodies. The following primary antibodies were used: anti-phospho-ACC S79 (Rabbit, 1:1000, #3661S, Cell signaling, Danvers, MA, USA), anti-AMPK (Rabbit, 1:1000, #2532, Cell signaling), anti-phospho-AMPK T172 (Rabbit, 1:1000, #2531, Cell signaling), Anti-ATP5I (Rabbit, 1:500, #16483-1-AP, Proteintech, Rosemont, IL, USA), anti OXPHOS cocktail human (Mouse, 1:750, #ab110411, Abcam, Toronto, ON, Canada), anti-F1-ATPase β-subunit (Mouse, 1:1000, #MABS1304, Sigma-Aldrich), anti-OSCP (Mouse, 1:1000, #ab110276, Abcam), anti-ATP5L (g) (Rabbit, 1:1000, #ab126181, Abcam), anti-GAPDH (Goat, 1:3000, #NB300-320, Novus Biologicals, Centennial, CO, USA), anti-α-Tubuline (Mouse, #T6074, Sigma Aldrich) and anti-β-Actin (Mouse, 1:10000, #3700, Cell signaling). The following secondary antibodies were used: anti-rabbit IgG conjugated to HRP (Goat, 1:3000, #170-6515, Bio-Rad), anti-mouse IgG conjugated to HRP (Goat, 1:3000, #170-6516, Bio-Rad) and anti-goat IgG conjugated to HRP (donkey, #Sc-2020, Santa Cruz Biotechnology, Dallas, TX, USA).

### Immunofluorescence

For immunofluorescence experiments, 50,000 - 100,000 cells were seeded on 1.5 mm thickness coverslips and were incubated for 48 h. As previously described^84^, cells were washed twice with ice cold PBS, then were fixed with a 4% paraformaldehyde solution for 10 min at 4 °C and washed 3x 10 min with a solution of PBS + 0.1 M glycine at rt with agitation. At this moment coverslips were sometime stored in PBS + 0.2 % Sodium Azide at 4°C until use. If stored, 3x 10 min washing steps with PBS were performed to remove the azide. Cells were then permeabilized with PBS + 0.2 % Triton X-100 and 3% Bovine Serum Albumin (BSA) for 5 min at rt. Then, cells were washed in a blocking solution (3 % BSA in PBS pH 7.4) 3x 10 min at rt with agitation and were then incubated with an appropriate diluted solution of primary antibody (in 3% BSA, PBS pH 7.4) o/n at 4°C in a humidified chamber. After primary antibody incubation, cells were washed in blocking solution 3x 10 min and were incubated with an appropriate diluted solution (3% BSA, PBS pH 7.4) of secondary AlexaFluor antibody for 1h at rt in the dark. Cells were then washed with PBS 3x 10 min and coverslips were mounted on glass coverslip in Vectashield mounting media containing DAPI (H-1200-10, Vector Laboratories, Newark, CA, USA) and were sealed off with nail polish and kept for a minimum of 24 hours at 4°C. Images were collected with the Zeiss Axio Imager Z2 upright microscope equipped with a CoolSNAP FX camera (Photometrics), Axiocam camera and ZEN 2 blue edition software and were analyzed using ImageJ software. Colocalization was assessed using the Job Plot profile functionality of ImageJ by determining fluorescence intensity for each pixel of each channel across a line drawn. For quantifications and colocalizations, raw data were exported to Prism (10.2.2, GraphPad) software to generate final figures. The following primary antibodies were used: Anti-ATP5I (Rabbit, 1:100, #16483-1-AP, Proteintech) and TOMM20 (Mouse, 1:100, #sc-17764, Santa Cruz Biotechnology). The following secondary antibodies were used: anti-mouse AF488 (Goat, 1:2000, #A-11029, Invitrogen), anti-rabbit AF568 (Goat, 1:2000, #A-11011, Invitrogen) and Streptavidin AF488(#S11223, Invitrogen). For automated quantifications, all images were first deconvoluted to enhance signal clarity. Mitochondrial morphology and network were then analyzed using the Mitochondria Analyzer plugin for Fiji.^85^

### TMRE-Based Mitochondrial Membrane Potential Assay

Mitochondrial membrane potential was assessed by live-cell fluorescence imaging using the cationic, lipophilic dye tetramethylrhodamine ethyl ester (TMRE; #T669, ThermoFisher Scientific). Cells were incubated with 100 nM TMRE in complete DMEM (with serum, without phenol red) for 30 minutes at 37 °C in a humidified CO incubator. For depolarization controls, cells were pretreated with 10 µM FCCP for 30 minutes prior to TMRE addition. Immediately before imaging, cells were washed twice with DMEM without phenol red (#319-080 CL, Wisent) to remove excess dye. Z-stack images were acquired from live cells using a Zeiss Axio-Observer Z1 spinning disk confocal microscope (63× objective, 12 optical slices). Three-dimensional reconstruction and quantitative analysis were performed using Fiji (ImageJ). Mitochondria were segmented using a custom macro based on Otsu’s thresholding method, followed by region of interest (ROI)-based quantification of area, mean intensity, perimeter, and integrated density via the ROI Manager. To verify that changes in TMRE signal reflected alterations in membrane potential rather than mitochondrial content, cells were co-stained with 50 nM MitoTracker Green (#M46750, ThermoFisher Scientific) under the same conditions. Since MitoTracker Green accumulates in mitochondria independently of membrane potential, it served as a control for mitochondrial mass. Results from this control experiment are not shown.

### Mitochondrial protein isolation

Mitochondrial crude extracts were obtained using Mitochondria Isolation Kit for Cultured cells (ab110170, Abcam) according to manufacturer’s instructions. For each condition, 1x 15 cm cell culture dish of KP-4 cells was washed twice in ice-cold PBS, then the residual PBS was removed by aspiration. Cells were then scraped in ice-cold PBS and transferred in Eppendorf tubes. The following steps for mitochondrial isolation were performed as described in the manufacturer’s instructions. The pellet containing mitochondrial proteins was resuspended in lauryl maltoside buffer [1% lauryl maltoside, cOmplete-EDTA free Protease Inhibitor Cocktail (Roche, Laval, Qc, Canada), PhosSTOP (Roche), PBS pH 7.8]. The concentrations of mitochondrial proteins were determined using the bicinchoninic acid assay (BCA) (23225, Pierce). Of note, at the end of isolation, we obtained ∼ 0.7-1 mg of mitochondrial proteins per 15 cm dish.

### Mitochondrial protein for BN-PAGE

Mitochondria were isolated using a procedure based on the manufacturer’s protocol (Mitochondria Isolation Kit for Cultured Cells, #ab110170, Abcam), with optimized reagent volumes adapted for smaller-scale preparations. While the standard protocol recommends four 15-cm culture dishes per isolation to maximize yield, we obtained satisfactory results using only two 15-cm dishes, consistently recovering 500–1000 µg of mitochondrial protein. Briefly, cells were harvested by scraping in ice-cold 1X PBS and transferred into two pre-chilled 1.5 mL microcentrifuge tubes to balance subsequent centrifugations. Cells were pelleted by centrifugation at 1000 × g for 3 minutes at 4 °C. To improve membrane disruption, a freeze–thaw cycle was performed by incubating cell pellets for 10 minutes on dry ice–ethanol, followed by 10 minutes at 37 °C. Pellets were then resuspended in 500 µL of Reagent A and subjected to mechanical homogenization using a pre-chilled 2 mL glass Dounce homogenizer. Thirty strokes were applied to ensure sufficient cell disruption. The homogenate was centrifuged at 1000 × g for 10 minutes at 4 °C to remove nuclei and cell debris. The supernatant was transferred to a new tube, mixed with 500 µL of Reagent B, and the protocol was followed as per the manufacturer’s instructions for subsequent steps. The final mitochondrial pellet was resuspended in 250 µL of the final buffer. Two aliquots of 100 µL were reserved for downstream purification experiments, while a 50 µL aliquot was kept for BCA quantification and OXPHOS western blot quality control. All samples were snap-frozen and stored at −80 °C until use.

### Blue native-PAGE

As part of the adapted protocol^50^, mitochondria (100 µg of protein) were lysed on ice for 30 minutes in a digitonin buffer containing 1% (w/v) digitonin (#11024-24-1, Sigma-Aldrich), 50 mM potassium acetate, 30 mM HEPES-KOH, 10% (v/v) glycerol, and EDTA-free complete protease inhibitors, pH adjusted to 7.4. Lysates were clarified by centrifugation at maximum speed for 60 minutes at 4 °C. Coomassie Brilliant Blue G-250 (#1610406, Bio-Rad) was then added to a final concentration of 0.1% (w/v) prior to electrophoresis. Proteins were resolved on NativePAGE™ 3–12% Bis-Tris gels (#BN1001BOX, Thermo Fisher Scientific) using a discontinuous buffer system with an anode chamber (outer) and cathode chamber (inner). Electrophoresis was conducted at 4 °C for 16 hours. The 20X running buffer used to prepare both the anode and cathode buffers contained 50 mM Bis-Tris and 50 mM Tricine, pH 6.8. The anode buffer was prepared by dilution to 1X. The 20X cathode additive contained 0.4% (w/v) G-250. Two working cathode buffers were used sequentially: a dark buffer with 0.02% G-250 and a light buffer with 0.002% G-250, both diluted from the 20X stock. Electrophoresis was initiated with the dark cathode buffer and anode buffer at 100 V for 2 hours at 4 °C. The dark buffer was then replaced with the light buffer, and electrophoresis was continued for approximately 14 hours at 150 V at the same temperature. Proteins were transferred onto a PVDF membrane pre-activated in methanol for 30 seconds, rinsed with water, and equilibrated for 5 minutes in a transfer buffer containing 25 mM Bicine, 25 mM Bis-Tris, and 1 mM EDTA, pH 7.2. The transfer was performed at 100 V for 1 hour and 30 minutes on ice. Following transfer, the membrane was incubated in 8% (v/v) acetic acid for 15 minutes, rinsed with water, briefly treated again with methanol for 30 seconds, rinsed once more with water, and then blocked according to the immunoblotting procedure described in this study. Immunodetection was carried out using an anti-ATP Synthase subunit β antibody (#MABS1304, Sigma-Aldrich). Signal quantification was performed with ImageJ software by measuring the area under the curve.

### Pull-down assay

For pull-down assay, 1 mg of mitochondrial proteins were incubated with either 1 mM of biotin functionalized biguanide (BFB) or biotin functionalized amine (BFA) in Eppendorf tubes for 3h at rt with mild agitation. In parallel, Dynabeads MyOne streptavidin C1 (65001, Thermo Fisher Scientific) were washed and resuspended at least three times in lauryl maltoside (LM) buffer [1% lauryl maltoside, cOmplete-EDTA free Protease Inhibitor Cocktail (Roche), PhosSTOP (Roche), PBS pH 7.8]. After incubation with biotinylated molecules, beads (0.5 mg Ill 50 μL) were added to mitochondrial lysates and incubated for 30 min at 4°C with agitation. Beads were respectively recovered with a magnet, washed in LM buffer 3x 30 min at 4°C. Elution of bound proteins from beads in BFB condition was performed by adding 30 μL of 50 mM metformin in LM buffer. After an incubation time of 30 min at rt, beads were vortexed, pelleted with a magnet and the supernatant was collected in a new tube. Proteins were then denatured by adding 30 μL of 6x SDS-loading buffer (30% glycerol, 10% SDS, 1% bromophenol blue, 15% β-mercaptoethanol, 0.5 M Tris-HCl pH 6.8) to the mixture and boiled 3x 5 min at 95°C. Elution of bound proteins for BFA condition and remaining proteins on beads for BFB condition were achieved by boiling and vortexing the corresponding beads in 60 μL of 6x SDS loading buffer 3x 5 min at 95°C. After denaturation of proteins, all samples were loaded into SDS–PAGE. The resulting gel was stained with Coomassie brilliant blue R-250 (#33445225GM, Thermo Fisher Scientific) then washed with a destaining solution (40% water, 50% methanol, 10% acetic acid). Gel segments were then excised, digested with trypsin and dried overnight in a cold trap (Labconco, Kansas City, MO, USA). Three samples for each elution condition were prepared in two technical replicates and were then analyzed by the Mass spectrometry platform at IRIC (Institute for Research on Immunology and Cancer). It’s worth mentioning that for pull-down validation, same experiments were performed using western-blot analysis with anti ATP5I antibody after protein denaturation steps (see immunoblotting section).

### Proteomics

Samples were reconstituted in 50 mM ammonium bicarbonate urea 8M vortexed and further diluted to 50 mM ammonium bicarbonate urea 1M with 10 mM TCEP [Tris(2-carboxyethyl)phosphine hydrochloride; Thermo Fisher Scientific], and vortexed for 1 h at 37°C. Chloroacetamide (Sigma-Aldrich) was added for alkylation to a final concentration of 55 mM. Samples were vortexed for another hour at 37°C. Trypsin 1 μg) was added, and digestion was performed for 8 h at 37°C. Samples were dried down and solubilized in 5% ACN-4% formic acid (FA). The samples were loaded on a 1.5 μl pre-column (Optimize Technologies, Oregon City, OR). Peptides were separated on a home-made reversed-phase column (150-μm i.d. by 200 mm) with a 56-min gradient from 10 to 30% ACN-0.2% FA and a 600-nl/min flow rate on a Ultimate 3000 connected to a Q-Exactive Plus (Thermo Fisher Scientific, San Jose, CA). Each full MS spectrum acquired at a resolution of 60,000 was followed by tandem-MS (MS-MS) spectra acquisition on the most abundant multiply charged precursor ions for 3s. Tandem-MS experiments were performed using higher energy collision dissociation (HCD) at a collision energy of 30%. The data were processed using PEAKS 7 (Bioinformatics Solutions, Waterloo, ON) and a Uniprot database. Mass tolerances on precursor and fragment ions were 10 ppm and 0.01 Da, respectively. Fixed modification was carbamidomethyl (C). Variable selected posttranslational modifications were acetylation (N-ter), oxidation (M), deamidation (NQ), phosphorylation (STY).

### Constructs

For recombinant protein expression in bacteria, full-length of human ATP5I sequence was PCR-amplified and cloned using BamH1/EcoRI restriction sites into a pET-TEV vector (Addgene, Watertown, MA, USA) for the expression of a N-terminal 6x-His-tagged protein. This plasmid was obtained as a kind gift from Dr. J.G. Omichinski (University of Montreal, Montreal). For lentiviral mediated transduction in KP-4 cells, annealed and phosphorylated oligos for two RNA guides (sgRNAs) targeting two regions of ATP5I’s cDNA (sgATP51 #1 and sgATP5I #2) and one RNA guide targeting GFP (sgGFP) sequence were subcloned into BsmBI restriction site of lentiCRISPRv2 plasmid (#52961, Addgene). For re-expression of ATP5I in control (sgGFP) and ATP5I knockout (KO) (sgATP5I#1 and #2) cells, full-length human ATP5I sequence with modified codons (same amino acid but different nucleic acid sequence) was generated by PCR site-directed mutagenesis and cloned using BamH1/EcoRI restriction sites into MSCV retroviral vector. All constructs were confirmed by DNA sequencing.

Constructs primers used:

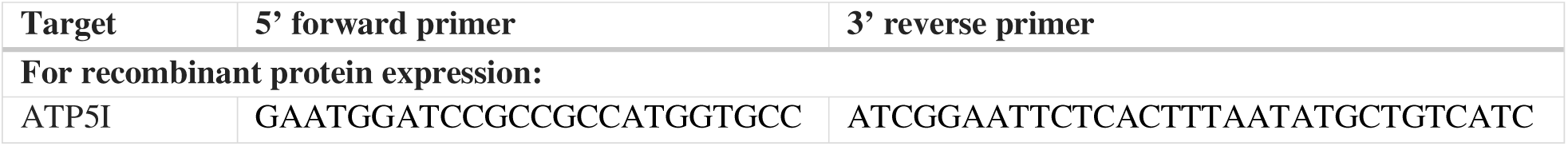
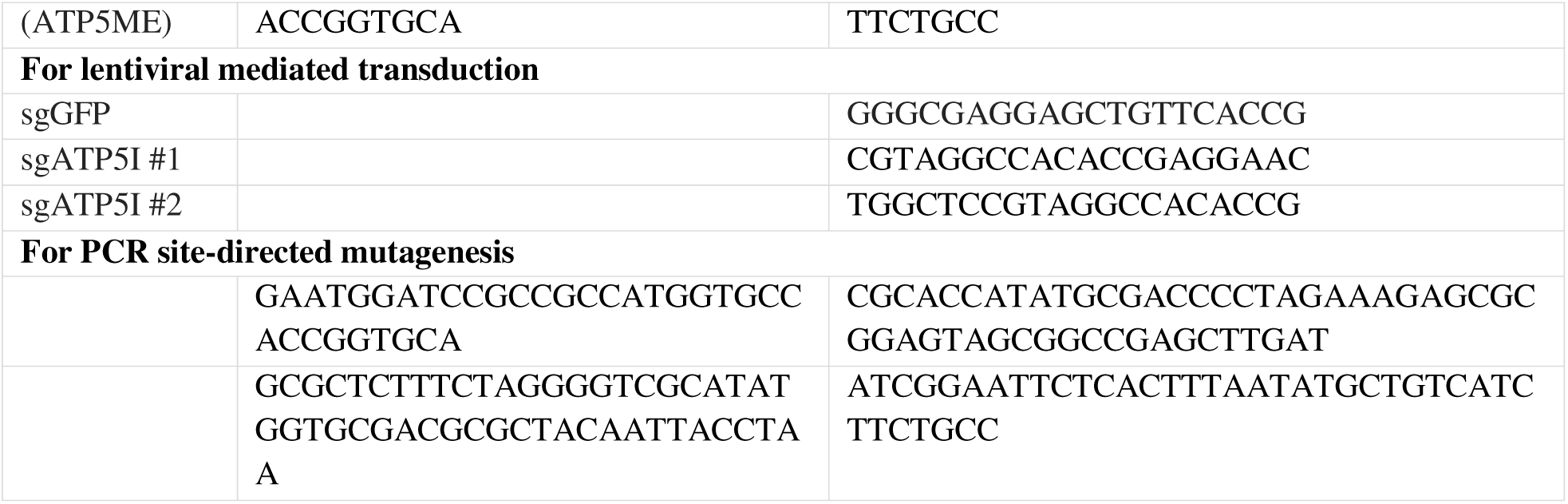

### Protein expression and purification

N-terminal 6x-His-tagged ATP5I was expressed in Rosetta E. coli BL21 competent cells (#70954, Addgene) for the expression of eukaryotic proteins that contain rare codons. To overexpress recombinant protein as suggested^86^, cells were grown at 37°C in terrific broth (TB) medium supplemented with 1% of ethanol (v/v), 100 mg/mL ampicillin and 50 mg/mL chloramphenicol to an OD_600nm_ of 0.8. Expression was induced for 4h at 30°C with 1 mM isopropyl β-D-1-thiogalactopyranoside then cells were harvested by centrifugation (15 min, 5000 rpm at 4°C). Bacterial pellets were then solubilized in buffer A (6 M guanidine, 500 mM NaCl, 20 mM NaH_2_PO_4_ pH 8) for 30 min at 4°C and lysed by sonication. Cell lysate was then harvested by centrifugation (30 min, 13 000 rpm at 4°C) and after filtration (0.45μm) the supernatant was loaded in an immobilized metal ion affinity chromatography column (HisTrap FF, Cytiva) with buffer A. After removing non-specifically bound molecules (flow through), the column was washed with buffer B (8M urea, 500 mM NaCl, 20 mM NaH_2_PO_4_ pH 8) then with buffer C (20 mM imidazole, 500 mM NaCl, 20 mM NaH_2_PO_4_ pH 8) until UV absorbance returned to baseline. After washing, the column was eluted with buffer D (500 mM imidazole, 500 mM NaCl, 20 mM NaH_2_PO_4_ pH 8) and the resulting eluate was loaded in a desalting column (HiPrep 26/10, Cytiva) with a PBS containing 125 mM NaCl, 5 mM DTT, PBS pH 7.4) and concentrated in a 3 kDa centrifugal filter (Amicon) to 1.5 mg/mL. Of note, all purification steps were done with AKTA pure chromatography system.

### Surface plasmon resonance (SPR)

Experiments were performed at 25°C with a portable SPR (P4SPR) device from JF. Masson’s laboratory (University of Montreal) (WO/2010/130045, Affinité instruments) using a compatible prism coated with a 200 nm streptavidin (SA) derivatized carboxymethyldextran hydrogel (Xantec bioanalytics) for immobilisation of biotinylated ligands. For each solution, a volume of 300 μL was injected in the flow cells comprising a multifluidic channel cell and a reference cell. After having stabilized the baseline (Ill30 min) with running buffer (125 mM NaCl, 5 mM DTT, PBS pH 7.4), biotin functionalized biguanide chloride salt was immobilized on SA sensor chip at 125 μM (concentration experimentally determined to saturate streptavidin sites). Then, the chip was washed with running buffer until stabilising the baseline. After that, several concentrations of purified recombinant ATP5I were injected until saturation of the signal. Of note, between each concentration added, washing steps were performed with running buffer to remove non-specifically bound molecules. Finally, the surface was regenerated with urea (2M) prepared in milli-Q-water. It’s worth nothing that due to the low dissociation constant of biotin/streptavidin interaction, biotinylated ligands are almost irreversibly bound that make it difficult to regenerate the chip. The sensorgrams were analysed with P4SPR Control v2.022 software and final figures were exported to Prism (10.2.2, GraphPad) using a pseudo-first order association kinetics equation for each concentration. For binding affinity curve, each steady state for each concentration were exported to Prism (10.2.2, GraphPad) software to generate the figure using a one site-specific binding equation. Of note, compounds were dissolved in the running buffer.

### Generation of stable ATP5I knockout cells

ATP5I Knock out in KP-4 cell line was done with lentiviral CRISPR/Cas9 technology and sgRNAs against ATP5I or a control sgRNA (see construct section). First, 5 x 10^6^ 293T cells were seeded in 10 cm plates and grown for 16 hours. They were then transfected using the calcium-phosphate precipitation method as previously described^87^ with 3 μg of lentiCRISPRv2 plasmid vector, 2 μg of the *pCMV-dR8.2* plasmid and 1 μg of the *VSV-G* envelope protein expression plasmid (both kind gift of Dr. N. Bardeesy). After 16 h, 10 mM sodium butyrate (Sigma Aldrich) was added for a minimum of 6 h, and then the medium was changed. In parallel, target cells were seeded to obtain 60% confluence the following day. Media from the transfected cells were collected the following day, filtered through a 0.45 μm filter. This viral soup was supplemented with 4 μg/mL polybrene (Sigma Aldrich) and 10% fresh media before being placed on KP-4 receiving cells. Viral soup was removed 24 hours later for fresh DMEM cell culture media (see cell culture section) supplemented with 100 μg.mL^-1^ sodium pyruvate (#600-110-EL, Wisent) and 50 μg.mL^-1^ uridine (#URD222.10, Bioshop) to help them growth in case mitochondrial functions were affected^88^. KP-4 infected cells were selected 6 hours after media change with 2ug/mL puromycin (#400-160-EM, Wisent). After two weeks in culture to ensure time alterations in the targeted regions with CRISP/CAS9, cells were replated highly diluted to isolate clones of cells. Many clones were grown and characterized by ATP5I immunoblots to select proper ATP5I knock-out.

### Retroviral infections

ATP5I re-expression was performed by retroviral-mediated expression of wild-type ATP5I in control (sgGFP) and ATP5I KO cells (sgATP5I #1 and #2) with a MSCV-ATP5I vector (see constructs).

For retroviral infections, 5 x 10^6^ Phoenix-Ampho packaging cells were seeded in 10 cm cell culture dishes and grown for 24 hours. Then, cells were transfected with 20 μg of MSCV-ATP5I vector and 10 μg of Helper envelope protein expression plasmid using calcium phosphate method as previously described^87^. The next day, sodium butyrate was added at 10 mM for 6 hours and then fresh medium was added. In parallel, target cells were seeded to obtain 60-70% on confluence the day of infection. Two days following transfection, the viral soup from one 10 cm dish was filtered through a 0.45μm filter, supplemented with 4 μg.ml^−1^ polybrene (#H-9268, Sigma Aldrich) and 1/10 (v/v) of fresh media and then used to infect one 10 cm dish of target cells. After at least 12 h post infection, infected cells were seeded into a new dish. After at least 4 hours of adherence, infected cells were selected using 50 μg/mL hygromycin (#400-141-UG, Wisent). Of note, the selection was maintained for the length of the experiments.

### Quantitative PCR (qPCR)

RNA isolation and qPCR quantifications were performed as previously described^83^ using the following primers (see below, BioCorp, Pierrefonds, Qc, Canada). Mitochondrial to nuclear DNA ratio were performed similarly by qPCR on 30 ng of DNA purified from cultures KP-4 cells. For the isolation, one 6 cm plate of sub confluent KP-4 cells were rinsed twice in PBS, collected by scraping in 500μL of SNET lysis buffer (5 mM EDTA, 400 mM NaCl, 1% SDS, 400 ug/mL Proteinase K, 20mM Tris-HCL pH8.0 [Sigma Aldrich]) and incubated at 55°C overnight. Equal volume of phenol:chloroform (Sigma Aldrich) was added and left to agitate for 30 min at rt. After a 5-minute centrifugation (16,000g), the aqueous phase was transferred to a new tube and precipitated by an equal volume of isopropanol, mixed well and DNA was collected by centrifugation at maximum speed (16,000g) for 15 minutes at 4°C. Pellet was washed with 70% ethanol, air dried and resuspended in TE.

qPCR primers used:

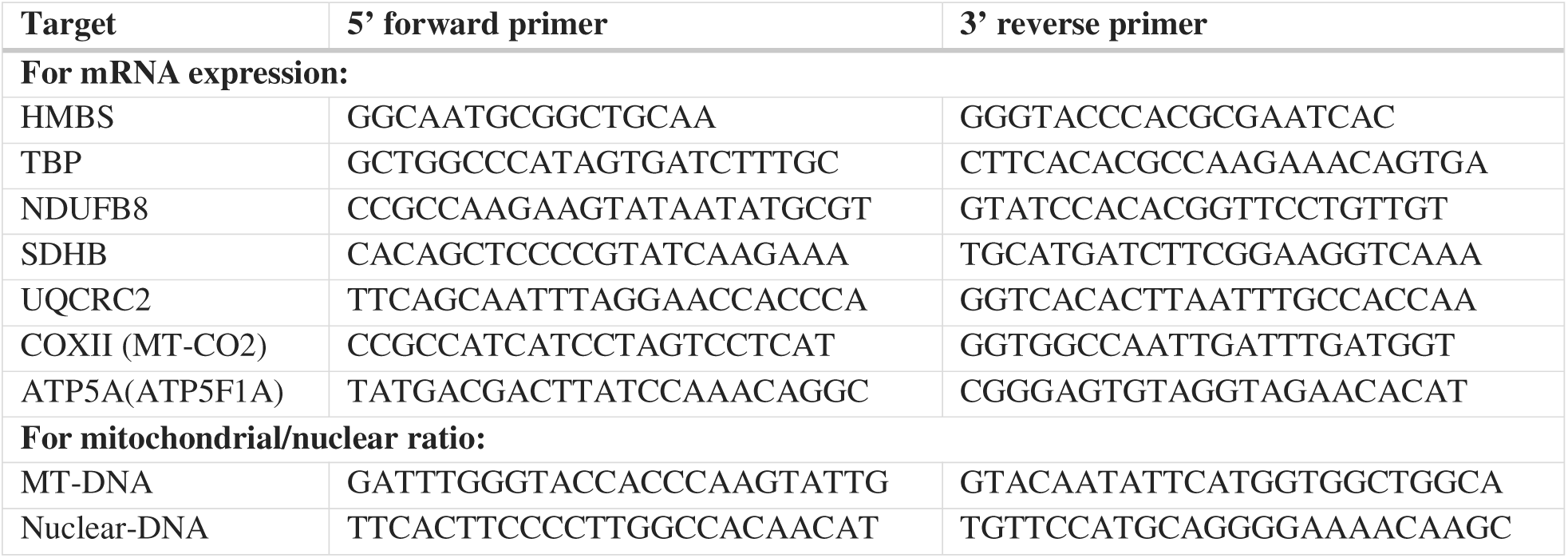

### NAD^+^/NADH quantification

For NAD^+^/NADH quantifications, 5×10^5^ cells were seeded in 10 cm cell culture dishes for technical duplicates and incubated for 48h. NAD^+^/NADH ratio were quantified using a fluorometric kit assay (#MAK460-1KT, Sigma Aldrich) following manufacturer’s instructions. For sample preparation, 9×10^5^ cells were pelleted and homogenized with corresponding extraction buffers to be in the linear range of the kit. Fluorescence intensities were measured at λ_Ex_ = 530 nm/λ_Em_ = 585 nm on SPARK 10 M (TECAN) and data were exported to Prism (10.2.2, GraphPad) software to generate final figures.

### Seahorse experiments

Seahorse XFe96 cell culture microplates (Agilent) were coated with 50 µg/mL of Poly-D-Lysine (#A3890401, Thermo Fisher Scientific) in D-PBS (#311-425-CL, Wisent) for 1 hour at room temperature and extensively washed with sterile water. Seahorse XFe96 sensor cartridges (Agilent) were calibrated according to the manufacturer’s protocol. Assay media was prepared as follows: 828 g DMEM without L-Glutamine, phenol red, sodium pyruvate, and sodium bicarbonate (#219-060-XK, Wisent), 29.2 mg L-glutamine (#AC386032500, Fisher Chemical), 450.4 mg D-glucose (#D-16-500, Fisher Chemical), 100 µL HEPES 1M (#330-050, Wisent), completed at 100 mL with Milli-Q water and adjusted pH at 7.2. Cells were plated in Poly-D-Lysine-coated microplates at a density of 40,000 cells/well in 100 µL of assay media. Following a 30-minute incubation at room temperature, 75 µL of assay media was added to each well, and the plate was transferred to an XFe96 analyzer. Typical runs were composed of the following steps: 1) baseline measurements for 18 minutes (3 cycles, each of 3-minute mixing and 3-minute measurement), 2) single drug injection (5 mM metformin, 100 µM phenformin, or sterile water as vehicle control), and 3) measurements over a period of 10.5 hours (63 cycles, each of 6-minute mixing, 1-minute pause, and 3-minute measurement). Oxygen consumption rates (OCR) and extracellular acidification rates (ECAR) were extracted from the Wave software (Agilent) and all time points were normalized to the first measurement. Data were then exported to Prism (10.2.2, GraphPad) software to generate final figures.

### Seahorse Bioenergetic Mito Stress Test

Cells were seeded in Poly-D-Lysine-coated XF96 microplates at a density of 40,000 cells per well in 100 µL of complete culture medium (see *Cell culture* in *Materials and Methods*). After a 1-hour incubation at room temperature to allow cell attachment, the medium was replaced with Seahorse assay medium described above by performing three gentle washing steps, leaving 50 µL per well. Cells were then treated with 125 µL of the appropriate condition: either metformin (5 mM), phenformin (100 µM), or sterile water as vehicle control. Plates were incubated for 3 hours at 37 °C in a non-CO incubator before being transferred to the Seahorse XFe96 analyzer. Typical runs consisted of a baseline measurement phase followed by sequential injections of mitochondrial inhibitors: oligomycin (10 µM), FCCP (3 µM), and a mix of rotenone and antimycin A (1 µM each). Each injection step included three measurement cycles (3-minute mixing and 3-minute measurement), for a total of 18 minutes per compound. Oxygen consumption rates (OCR) and extracellular acidification rates (ECAR) were obtained using the Wave software (Agilent).

### Crystal violet cell number assay

The crystal violet staining retention assay^89^ was used to estimate cell growth and viability by measuring absorbance at 590 nm in a microplate reader (SpectraMax 190 microplates, Molecular Devices). For the growth assay, 5,000 cells were seeded in 4 different 6-well cell culture dishes in technical triplicates and incubated for 4 different time points up to 6 days. Of note, media was replaced every 48 hours. At indicated times, cells were washed twice in PBS and fixed in 1% glutaraldehyde solution (PBS) for 10 min at rt. After fixation, cells were washed twice in PBS. Once all time points were collected, plates were stained in a 0.05 % crystal violet solution (PBS) for 30 min with agitation. After staining, plates were washed five times in water and were dried for 24 hours. The next day, crystal violet retained in cells were solubilized in 10 % acetic acid solution for 15 min with agitation and transferred into a 96 well plates for absorbance measurement. Data were then exported to Prism (10.2.2, GraphPad) software to generate final figures.

For the cell viability assay, 1,000 cells were seeded in 96 well plates in technical triplicates. After 24 hours, cells were treated at different concentrations with the corresponding drugs and were grown for 72 hours. Crystal violet staining was performed as described for the growth assay. The half minimal effective concentrations were obtained using a fit a curve with non-linear regression log(inhibitor) vs. response -variable slope (four parameters) from the corresponding dose response curves with Prism (10.2.2, GraphPad) software. Importantly, because growth rate was affected in ATP5I KO cells generating variabilities, cell viability assays comparing ATP5I KO with control cells were done in media supplemented with pyruvate and uridine (see generation of stable ATP5I KO cells section). Of note, drugs were dissolved in the corresponding media.

### ATP quantification

For ATP measurements, 3×10³ cells were seeded per well in a 96-well plate in technical duplicates. ATP levels (in picomoles) were determined using the ATP Determination Kit (#A22066, Invitrogen), according to the manufacturer’s instructions. Prior to incubation, cells were washed three times with DMEM without phenol red (#319-080 CL, Wisent). After a 15-minute incubation with firefly luciferase, luminescence was recorded at an emission wavelength of 560 nm (λ = 560 nm) using a SPARK 10M plate reader (TECAN). The data were subsequently exported to GraphPad Prism (v10.2.2) for figure generation.

### Chemogenomic CRISPR/Cas9 KO screen in NALM-6 cells

The Genome-wide pooled CRISPR/Cas9 KO screens made in the presence of chemicals inhibiting growth were performed by the ChemoGenix platform (IRIC, Université de Montréal; https://chemogenix.iric.ca/) as previously described^90^. Briefly, a NALM-6 clone bearing an integrated doxycycline-inducible Cas9 expression cassette generated by lentiviruses made from pCW-Cas9 (Addgene #50661) was transduced with the genome-wide KO EKO sgRNA library^90^ (278,754 different sgRNAs). After thawing the library from liquid N_2_ and letting it recover in 10% FBS RPMI for 1 day, KOs were induced for 7 days of culture with 2 ug/mL doxycycline. The pooled library was then split in different T-75 flasks (28×10^6^ cells per flask; a representation of 100 cells/sgRNA) in 70 mL at 4×10^5^ cells/mL. Cells were treated with 16 mM Metformin (using 1M stock solution prepared in media) or 70 nM rotenone (Sigma, R8875) or 2 µM oligomycin A (Tocris Bioscience, 4110) (both from 1000X stock solutions prepared in DMSO) for 8 days with monitoring of growth every 2 days, diluting back to 4×10^5^ cells/mL and adding more compounds to maintain same final concentration whenever cells reached 8×10^5^ cells/mL. Over that period, treated cells had respectively 1.47, 2.93 & 4.14 population doublings whereas no solvent or DMSO controls had about 7.5. Cells were collected, genomic DNA extracted using the Gentra Puregene kit according to manufacturer’s instructions (QIAGEN), and sgRNA sequences PCR-amplified as described^90^. SgRNA frequencies were obtained by next-generation sequencing (Illumina NextSeq 500). Reads were aligned using Bowtie2.2.5 in the forward direction only (norc option) with otherwise default parameters and total read counts per sgRNA tabulated. Context-dependent chemogenomic interaction scores were calculated from comparing sgRNA frequency changes against negative controls from using a modified version of the RANKS algorithm^90^ which uses guides targeting similarly essential genes as controls to distinguish condition-specific chemogenomic interactions from non-specific fitness/essentiality phenotypes. Raw read counts are available upon request directly from the ChemoGenix platform.

### Statistical analysis

Statistical analyses were performed using GraphPad Prism (10.2.2) software. Unpaired two-tailed Student’s *t*-test or Ordinary one-way ANOVA with compensation of multiple comparison with Sidak’s test were used to determine significance except for seahorse analysis where paired two-tailed Student’s *t*-test and RM one-way ANOVA were preferred. A value of p < 0.05 was considered significant. All specific statistical details can be found in the corresponding legends.

